# Structural Context Determines Docking Engine Performance: A Family-Stratified Benchmark of Six Engines

**DOI:** 10.64898/2026.08.11.744016

**Authors:** Kristoffer Alejo, Sarah Fisher, Tejaswan Kalluri, Balkrushna More, Hrishikesh Rajgure, Pritam Kumar Panda, Christopher Korban, Christian Chung

## Abstract

Molecular docking and co-folding engines are widely used to prioritize compounds for wet-lab validation, yet their accuracy is known to vary substantially across protein targets for reasons that remain only qualitatively understood. Here we benchmark six docking and co-folding engines (RevDock, DiffDock, Boltz2, AutoDock-GPU, rDock, and PandaDock) across 14 protein families, evaluating scoring power, ranking power, docking power, and physical validity. Rather than treating engine performance as protein-family-specific, we classify all 14 families into six mechanistic groups according to which of four scoring-function simplifications, rigid receptor, pairwise additivity, fixed point charges, and implicit solvent, is most severely stressed by that family’s binding site. This framework helps explain, rather than simply describe, where each engine succeeds or fails: RevDock’s CNN rescoring layer mitigates the pairwise additivity and fixed-charge limitations relative to physics-only scoring, achieving the highest overall pose accuracy (73.3% of poses ≤ 2.0 Å RMSD), while Boltz2’s sequence-based co-folding bypasses the rigid-receptor assumption and achieves comparable affinity correlation (mean Pearson *r* ≈ 0.60 for both engines). PandaDock, run with expanded conformational sampling, matches RevDock on pose accuracy (72.1% of poses ≤ 2.0 Å, lowest median RMSD at 0.96 Å) and exceeds AutoDock-GPU on affinity correlation (mean *r* = 0.460), indicating that the performance of a physics-based scoring function is limited as much by search adequacy as by the scoring function itself. These results suggest that engine selection for a docking or co-folding campaign should be guided less by an engine’s aggregate benchmark ranking and more by which of these four structural and physical characteristics dominate the target of interest.

## 1 Introduction

In 2016, the estimated cost of both the drug development process and go-to-market process skyrocketed by 145% in the past decade, and although the time to reach clinical trials has decreased, its success rate has dropped to only 12%^1^. This observation has shown how, regardless of the computational advances that have allowed the overall process to be expedited, the cost and rigor necessary to reach FDA approval have soared. Computational methods such as docking and co-folding have been identified as quintessential tools necessary for both the hit identification and lead optimization process, where the success is measured by the ability to produce meaningful hits while reducing the number of overall compounds necessary for wet-lab testing^2^. Prior literature has established molecular docking as an effective engine capable of large compound screening at low costs compared to HTS^3^, proposing a potential solution to reduce the overall cost of an end-to-end drug discovery campaign. While proving to be a cost-reducing measure, it places a question on the overall accuracy of the engine. Multiple robust docking algorithms are capable of accurately predicting a pose; however, it is suggested that there are major scoring function limitations associated with said engines^1^. While both docking and co-folding have similar functions in terms of pose geometry hypotheses and evaluation of pose plausibility, there are fundamental differences to note. Classical docking treats the receptor as rigid and therefore assumes the receptor to be ground truth. In comparison, co-folding utilizes the protein sequence and therefore pose evaluation is no longer just an evaluation of the ligand, but also the receptor structure as well^4^. These distinctions have direct implications for how each engine is evaluated, which we address in the following section.

Furthermore, the use of a rigid receptor and a defined binding site allows traditional docking to score a ligand conformation against a fixed protein structure. As a result, it is essential that a high-quality co-crystal structure is utilized to ensure the most biologically accurate results^4^. With co-folding, a protein sequence is taken with a SMILES string to predict the protein structure and ligand pose simultaneously. This distinction between docking and co-folding means that the six engines to be evaluated in this study are not doing the same thing. These engines differ in what inputs they require, what outputs they produce, and the claims they make about affinity. In this study, we aim to analyze the six docking and co-folding engines. RevDock, evaluated here, is our fork of GNINA^5;6^ and retains its Vina-style pose generation and CNN rescoring architecture; results reported for RevDock should therefore be read as characterising that architecture rather than an independently derived scoring function. Of these six engines, only Boltz2 and RevDock produce affinity-calibrated outputs, whereas the remaining four produce either physics-based binding energies or pose confidence scores that are not calibrated to an experimental affinity scale^5;7^. Only Boltz2 operates as a co-folding model, where the other five employ classical docking methods^7^. It is also worth noting that both DiffDock and Boltz2 require no binding site definitions, unlike the other four engines evaluated^7;8^.

The performance of each docking and co-folding engine cannot be measured holistically, as results may vary subsequently across protein families. A study benchmarking a series of co-folding models against both kinases and G protein-coupled receptors (GPCRs) was conducted, where they found a substantial difference in performance between the two protein families^9^. A protein family can be defined by tracking the lineage of a protein, where a group of proteins can be classified by a common ancestor. In addition, protein families are known to share similar structure, sequence, and function^10^. In regard to docking and co-folding, evaluating each engine on different protein families will allow us to derive insights on what physical structures or interactions each engine is able to capture effectively and which engine to utilize for any specific target. Table S1 (Supporting Information) characterizes all 14 protein families evaluated in this case study by structural hallmarks, binding site descriptions, dominant chemotypes, and the primary docking challenge for each family. Structural and functional differences across these families create fundamentally distinct docking challenges, motivating a family-stratified evaluation of each engine. Figure 1 summarises the aim and design of the study: how the benchmark set was assembled, which engines were compared and on what terms, the four axes along which they were evaluated, and the central finding that emerges.

**Figure 1:**
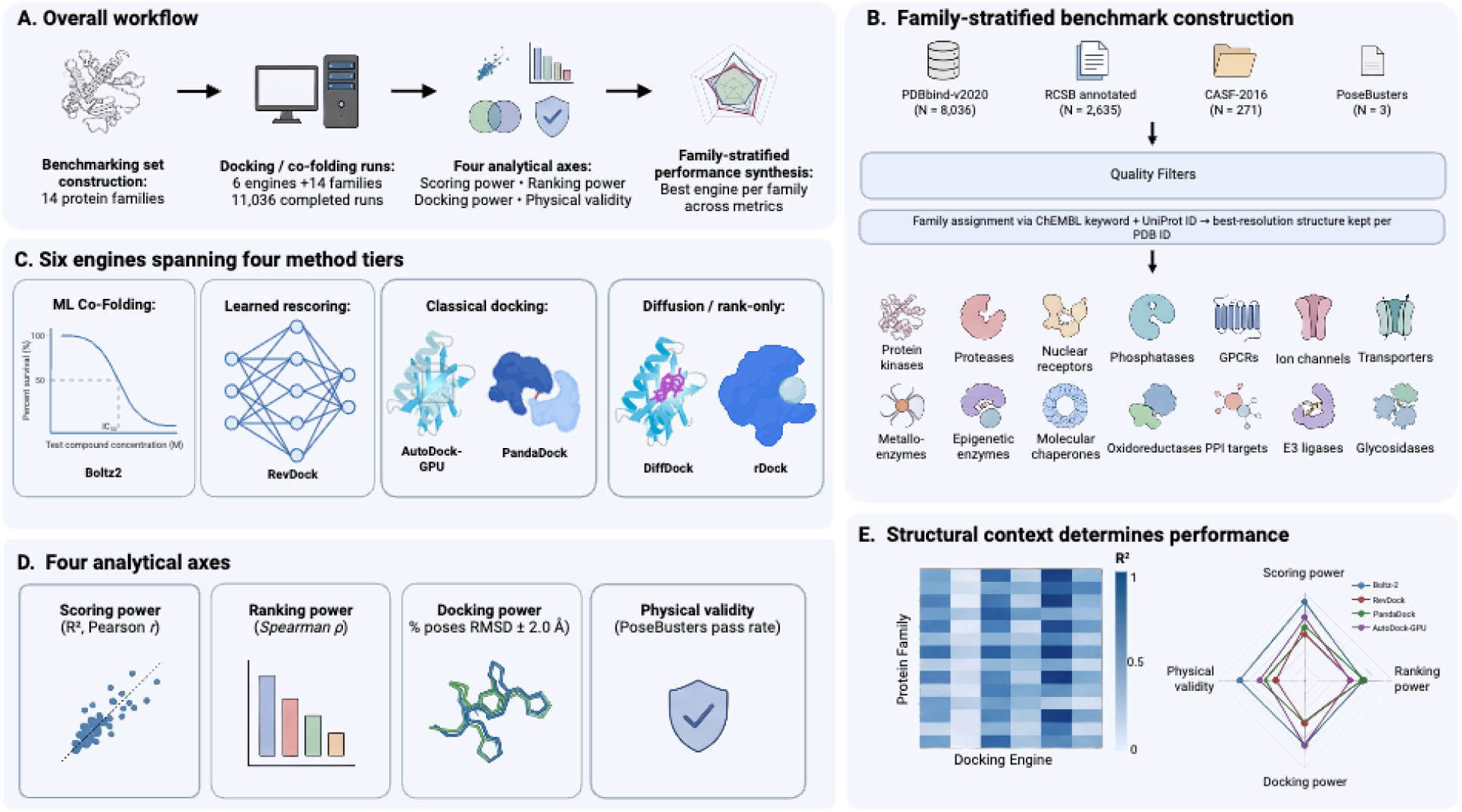
Overview of the study. (a) Workflow: benchmark construction, docking and co-folding runs, evaluation along four analytical axes, and family-stratified synthesis. (b) Benchmark construction from four source databases through quality and bioactivity filters to 14 protein families (11,036 of 12,420 attempted runs completed). (c) The six engines, grouped into four method tiers by what they consume and what their score means; only Boltz2 and RevDock are calibrated to an experimental affinity scale. (d) The four analytical axes: scoring power, ranking power, docking power and physical validity. (e) The central result, summarised as an *R*^2^ matrix over families × engines and as a four-axis performance profile per engine: performance is set by the structural context of the target family rather than by engine identity alone, and search budget contributes as much as the scoring function.

In this study, we aim to evaluate six docking/co-folding engines across 14 protein families to determine which engine performs best per family and overall. Each engine and protein family will be assessed along the following four analytical axes: scoring power (evaluate affinity correlation with *R*^2^ and Pearson r), ranking power (evaluate rank-order correlation with Spearman *ρ*), docking power (evaluate pose reproduction with RMSD), and physical validity assessment (evaluate ligand geometry using Posebusters pass rate). It is worth noting that not all engines are capable of being evaluated across all four axes as a result of the discrepancy between calibrated and uncalibrated outputs of each engine. As the experimental dataset utilized was binders and co-crystal structures only, no inactive or decoy compounds were available to conduct a screening power evaluation. Full methodological details for each analytical axis are described in the Methods section below.

## 2 Methods

To conduct this study, we utilized four primary databases. We retrieved 8036 structures from the PDBbind-v2020 set, 2635 structures from RCSB containing PDBbind affinity values, 271 structures from the CASF-2016 core set, and 3 structures from the PoseBusters benchmarking dataset. Targets were first identified via ChEMBL using family keywords, gene name, and protein classification (L1/L2/L3), where UniProt IDs were then batched into RCSB searches. From these databases, co-crystal structures were then selected using a bioactivity filter where each complex must contain at least one of the following records: IC50, Ki, Kd, EC50, or Inhibition. Quality filters were then applied, where only X-ray crystallography structures were utilized. In addition, each structure must have a resolution of ≤ 2.5 Å, R-free ≤ 0.30, a ligand with molecular weight between 100-800 Da, and ≥ 5 heavy atoms. Each PDB ID was then assigned to the best-resolution family, ensuring no duplicates across families. Furthermore, preprocessing for each ligand was conducted, where the following ligand types were excluded from the study: ions, cryoprotectants, buffer components, Fe-S clusters, crystallographic sugars, modified residues, and unknowns/water. It is worth noting that 54% of matched affinity entries were Ki or IC50 values rather than strict Kd values. This introduces systematic noise for engines whose scoring functions were calibrated to Kd. All structures were then classified into 14 protein families. In total, 11,036 of 12,420 attempted docking runs completed successfully, with failure rates varying from 0.1% for rDock to approximately 20% for AutoDock-GPU.

For all co-crystal structures, each complex was then prepared in the input formats required by each engine for downstream docking/co-folding runs. For RevDock and AutoDock-GPU, both the receptor and the ligand were stored as individual PDBQT files via Meeko. Where Meeko receptor conversion failed due to non-standard residues, a fallback pipeline was applied using Open Babel for protonation at pH 7.4 followed by AutoDockTools for receptor preparation. For DiffDock, PandaDock, Boltz2, and rDock, the receptor was stored as the raw PDB file, where the ligand was stored as an SDF for DiffDock, PandaDock, and rDock. Boltz2 stored the ligand as a SMILES string, which was extracted from the manifest or derived via RDKit MolToSmiles on the crystal SDF. For multi-fragment SDFs, the largest fragment by heavy atom count was selected as the docking ligand. For RevDock, AutoDock-GPU, and PandaDock, a docking box was established with center coordinates at the mean of crystal ligand heavy-atom coordinates for each co-crystal structure. The box size was defined as a cube, where the side length was determined by the largest ligand extent across any single axis plus a 10 Å buffer, rounded up to the nearest even integer. For AutoDock-GPU, grid maps were computed using AutoGrid4 at a grid spacing of 0.375 Å. For DiffDock and Boltz2, no grid box needed to be established. rDock established a ligand cavity where the cavity was defined with a 6.0 Å radius centered on the crystal ligand centroid, a grid step of 0.5 Å, and a maximum of one cavity per structure.

Following protein and ligand preparation, all inputs were passed through their respective engine, where for each engine, a pose prediction as well as pose scoring metrics were returned. Boltz2 returned an affinity prediction value measured in ln(IC50), an affinity probability score, and an overall confidence score. RevDock returned a CNN affinity score measured in pKd, CNN score, and minimized affinity score measured in kcal/mol. DiffDock did not return an affinity score, but rather a pose confidence score. PandaDock returned a best pose energy score measured in kcal/mol. AutoDock-GPU measured the best binding energy across 20 runs in kcal/mol. rDock used a dimensionless score where better pose quality was assessed with lower scores. As rDock scores are negative with lower values indicating better binding, scores were negated before correlation analysis to ensure positive Spearman *ρ* reflects correct ranking direction. It is worth noting that native scores are not directly comparable across engines as they differ in units, scale, and directionality; all cross-engine comparisons were therefore conducted using derived metrics rather than native scores directly.

Scoring and ranking power were then both assessed using affinity correlation, where scoring power was measured using both *R*^2^ and Pearson r, and ranking power was measured using Spearman *ρ*, where a two-sided p-value was computed for each Spearman *ρ*. Engines were assigned to tiers based on whether their output was calibrated to an experimental affinity scale, determining which correlation metrics were computed. Experimental ground truth affinity values were established as −*log*_10_(*affinity*), equivalent to pKd, pKi, and pIC50, where unit conversions were then performed to make an equivalent comparison for each engine. Correlations were computed within each protein family and engine group, with a minimum of five structures required per group for inclusion. For Boltz2, affinity correlation was calculated comparing predicted ln(IC50) against experimental ln(IC50). Experimental −*log*_10_(*affinity*) was converted to ln(IC50) with the following equation:

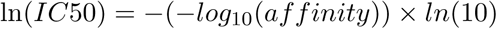

For RevDock, predicted CNN affinity in pKd was correlated against experimental −*log*_10_(*affinity*), equivalent to pKd. PandaDock and AutoDock-GPU took the top1_score in ΔG and correlated it against experimental ΔG, where −*log*_10_(*affinity*) was converted as follows:

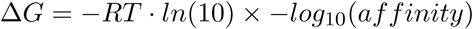

DiffDock and rDock correlated their respective scores against −*log*_10_(*affinity*). It is worth noting that DiffDock and rDock were used only for rank ordering, only measuring Spearman *ρ*, and not *R*^2^ and Pearson r. Boltz2, RevDock, PandaDock, and AutoDock-GPU were assessed with all three metrics. Within each protein family and engine group, predicted and experimental affinity values were Z-score normalized before visualization, enabling cross-engine visual comparison despite differing unit scales. An ordinary least squares regression line (OLS) was fitted per protein family and overlaid on each calibration plot alongside a y = x ideal predictor line.

RMSD analysis was performed on every docked structure for which a co-crystal reference was available, comprising 6,368 crystal ligand SDF files and 6,368 crystal protein PDB files and yielding 11,024 pose–crystal pairs across 2,064 structures. An earlier version of this analysis was restricted to a 1,053-structure subset; per-family conclusions differ between the two and only the complete set is reported here. Docking power was then assessed using RMSD analysis. For non-Boltz2 engines (e.g., RevDock, DiffDock, PandaDock, AutoDock-GPU, and rDock), both the predicted pose and crystal ligand SDF files were loaded with RDKit. Because some SDF files contained multiple disconnected fragments (e.g., salt counterions, co-crystallised solvent, etc.), only the largest connected fragment defined by heavy atom count was kept for both the predicted and crystal molecules, preventing the RMSD calculations from failing on disconnected graphs. Atom count validation was then conducted, ensuring the predicted and experimental structures contained an equal number of atoms; structures with a mismatch were excluded from RMSD calculations. RMSD was computed over heavy atoms without superposition, using a Hungarian assignment on the inter-atomic distance matrix to resolve the atom correspondence, so that molecular symmetry does not inflate the value through mismatched equivalent atoms. We note that this assignment is not element-constrained and may therefore pair atoms of different types in rare cases; a symmetry-corrected graph-automorphism RMSD^11^ would be marginally more conservative.

Boltz2 outputs a single PDB file containing both the predicted protein (chain A) and the predicted ligand (chain B). The ligand coordinates were then pulled directly from the raw PDB by selecting ATOM records assigned to chain B. The predicted protein was then aligned to the crystal protein, where the C*α* backbone atoms of the predicted protein were then aligned onto the C*α* backbone atoms of the crystal protein using the Kabsch algorithm, which aims to find the optimal rotation and translation that minimizes the distance between the two backbones. Where the predicted and crystal protein chains differed in length, both N-terminal and C-terminal trims were attempted, and the trim yielding the lower backbone RMSD was selected. The same transformation was then applied to the ligand coordinates extracted in the previous step. The Hungarian algorithm was then applied to find the optimal atom-to-atom assignment, as Boltz2’s PDB output did not guarantee that the atoms that are listed are in the same order as the crystal SDF. Atom pairings between elements of different types were penalized with a distance of 10^6^ Å^2^ to prevent chemically invalid assignments. RMSD was then computed over these matched pairs. It is worth noting that the retrieved subset is unevenly distributed across protein families, with Glycosidases having an insufficient sample size of (N = 12). Per-family results for Glycosidases were treated as preliminary. Docking power assessment was conducted by calculating the percentage of poses with RMSD ≤ 2.0 Å, with median RMSD reported as a secondary metric.

Physical validity testing was then conducted with PoseBusters^12^. PoseBusters is a rule-based cheminformatics validation tool that aims to validate the chemical and physical plausibility of a ligand, using a series of checks segmented into these three main subcategories: chemical consistency, intramolecular geometry, and energy ratio. The results of a PoseBusters validation are binary, where a pass is only assigned if all checks have passed. PoseBusters was conducted on all successful docking runs across all six engines and 14 protein families, totaling 11,036 pose-engine pairs. Only runs that produced a valid output pose file were included; failed docking runs were already excluded at the scoring stage. Before analysis, each predicted pose for SDF-based engines (e.g., RevDock, DiffDock, PandaDock, AutoDock-GPU, rDock) was loaded with RDKit, using a sanitisation fallback that retains molecules with valence violations so that they are evaluated by PoseBusters rather than silently dropped from the denominator. Boltz2 outputs a full protein-ligand complex compared to a ligand-only SDF file like the other engines. To extract the ligand, chain B was selected from the predicted complex, which yields correct coordinates but, because PDB records carry no bond orders, an all-single-bonded molecule. Bond orders were therefore restored by matching the extracted ligand against the input SMILES that Boltz2 was given, using RDKit AssignBondOrdersFromTemplate. This step is necessary rather than cosmetic: without it, PoseBusters’ intramolecular geometry and energy-ratio checks fail on essentially every aromatic ligand for reasons of representation rather than pose quality, and the Boltz2 pass rate is depressed from 94.8% to 40.6%. PoseBusters v0.6 was then run in mol mode on all preprocessed ligand molecules, where each pose received a binary pass flag. A five-second per molecule timeout was enforced; the 7 of 11,036 poses that timed out were excluded from pass rate calculations.

## 3 Results

Affinity correlation analysis was performed across all 14 protein families for four of the six engines: Boltz2, RevDock, AutoDock-GPU, and PandaDock. rDock and DiffDock were excluded from the scoring power assessment as their respective outputs of an empirical dimensionless score and a pose confidence score are not calibrated to an experimental affinity scale. Results show Boltz2 with a mean r = 0.604, *R*^2^ = 0.391, *ρ* = 0.571, and significant at p < 0.05 in 12 of the 14 families assessed. RevDock was assessed with a mean r = 0.605, *R*^2^ = 0.396, *ρ* = 0.566, and was significant in 12 of the 14 families. PandaDock, rerun with expanded conformational sampling (v4.1.1), was assessed with a mean r = 0.460, *R*^2^ = 0.246, *ρ* = 0.451, and was significant in 12 of the 14 families. AutoDock-GPU was assessed with a mean r = 0.403, *R*^2^ = 0.196, *ρ* = 0.403, and was significant in 12 of the 14 families assessed. *R*^2^ and Pearson r were not assessed for DiffDock. A mean *ρ* = –0.050 and significance in 6 out of 14 protein families was observed only as an exploratory rank-ordering analysis. The same applies to rDock where a mean *ρ* = 0.006 with significance in 4 out of 14 protein families was observed only as an exploratory rank-ordering analysis as well. Boltz2 performed best with PPI targets (*ρ* = 0.837), Metalloenzymes (*ρ* = 0.793), and Phosphatases (*ρ* = 0.744). Boltz2 performance showed the weakest performance on GPCRs (*ρ* = 0.305) and Transporters (*ρ* = 0.220) (Figure S1A). RevDock peaked on PPI targets (*ρ* = 0.871), Phosphatases (*ρ* = 0.814), and Protein kinases (*ρ* = 0.724), with weakest performance on Nuclear receptors (*ρ* = 0.379) and Transporters (*ρ* = 0.190) (Figure S1B). AutoDock-GPU performed best in PPI targets (*ρ* = 0.751) and Proteases (*ρ* = 0.590). PandaDock peaked on E3 ligases (*ρ* = 0.734, the best of any engine for that family), Phosphatases (*ρ* = 0.680), and PPI targets (*ρ* = 0.642), with weakest performance on Transporters (*ρ* = 0.032) and Nuclear receptors (*ρ* = 0.224) (Figure S1C).

**Figure 2:**
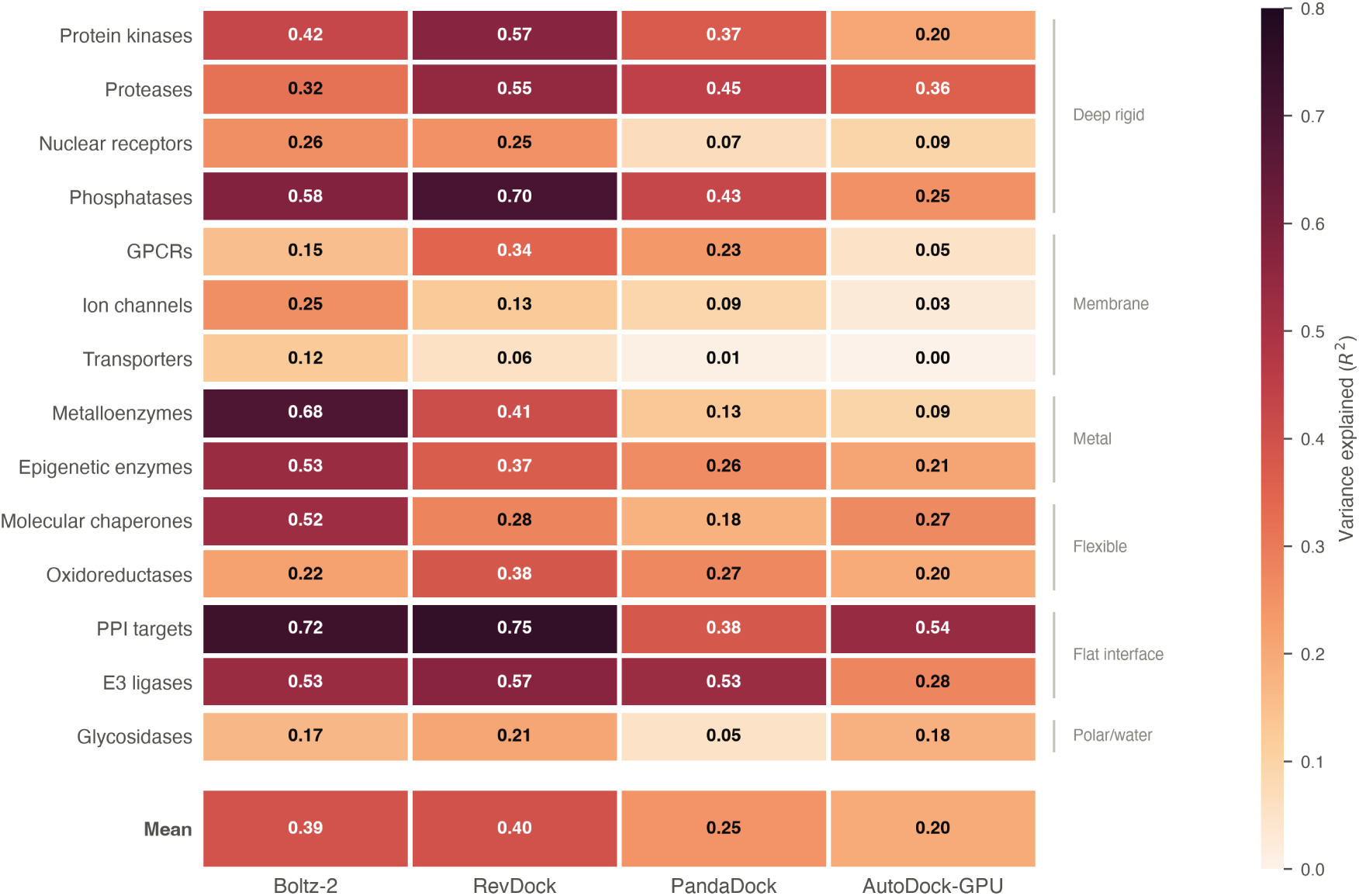
Scoring power by engine and target family. Cell values are *R*^2^ between predicted and experimental binding affinity, computed within each family. DiffDock and rDock are omitted because their outputs are not calibrated to an affinity scale. Right-hand brackets group families by the scoring-function simplification they most stress.

Boltz2 (Figure S1A) and RevDock (Figure S1B) panels show regression lines clustered around y = x for most families with visible positive slopes across all 14 families. The PandaDock panel now shows consistent positive slopes across 13 of 14 families (Figure S1C), replacing the near-flat, slope-free pattern obtained under default sampling. The AutoDock-GPU panel showed positive slopes but noticeably shallower fans compared to Boltz2 and RevDock (Figure S1D). PPI targets (Figure S2J), Phosphatases (Figure S2K), and Protein kinases (Figure S2M) showed the tightest regression alignment across both Boltz2 and RevDock. Transporters (Figure S2N) and GPCRs (Figure S2C) show the flattest lines.

**Table 1:** Affinity correlation results across 14 protein families for four docking engines. Pearson *r* and *R*^2^ (scoring power) were computed for Tier 1–3 engines only. Spearman *ρ* (ranking power) was computed for all tiers. DiffDock and rDock are excluded as Tier 4 engines whose outputs are not calibrated to an experimental affinity scale. Bold values indicate the best-performing engine per family per metric.

| Protein Family | Pearson $r$ | | | | $R^2$ | | | | Spearman $\rho$ | | | |
| --- | --- | --- | --- | --- | --- | --- | --- | --- | --- | --- | --- | --- |
|  | Boltz2 | RevDock | PandaDock | AutoDock-GPU | Boltz2 | RevDock | PandaDock | AutoDock-GPU | Boltz2 | RevDock | PandaDock | AutoDock-GPU |
| E3 ligases | 0.727 | **0.754** | 0.726 | 0.527 | 0.529 | **0.568** | 0.527 | 0.278 | 0.701 | 0.669 | **0.734** | 0.513 |
| Epigenetic enzymes | **0.727** | 0.606 | 0.511 | 0.459 | **0.529** | 0.368 | 0.261 | 0.210 | **0.729** | 0.612 | 0.528 | 0.485 |
| GPCRs | 0.389 | **0.587** | 0.485 | 0.228 | 0.151 | **0.345** | 0.235 | 0.052 | 0.305 | 0.416 | **0.431** | 0.088 |
| Glycosidases | 0.408 | **0.454** | 0.224 | 0.427 | 0.166 | **0.207** | 0.050 | 0.182 | 0.389 | **0.477** | 0.258 | 0.455 |
| Ion channels | **0.504** | 0.358 | 0.305 | 0.160 | **0.254** | 0.128 | 0.093 | 0.025 | **0.462** | 0.406 | 0.298 | 0.237 |
| Metalloenzymes | **0.826** | 0.639 | 0.359 | 0.306 | **0.682** | 0.409 | 0.129 | 0.094 | **0.793** | 0.558 | 0.368 | 0.323 |
| Molecular chaperones | **0.722** | 0.529 | 0.425 | 0.516 | **0.522** | 0.280 | 0.180 | 0.266 | **0.674** | 0.493 | 0.389 | 0.510 |
| Nuclear receptors | **0.505** | 0.501 | 0.259 | 0.306 | **0.255** | 0.251 | 0.067 | 0.094 | **0.497** | 0.379 | 0.224 | 0.306 |
| Oxidoreductases | 0.467 | **0.618** | 0.519 | 0.443 | 0.218 | **0.382** | 0.269 | 0.196 | 0.484 | **0.610** | 0.484 | 0.451 |
| PPI targets | 0.849 | **0.864** | 0.617 | 0.732 | 0.720 | **0.746** | 0.381 | 0.535 | 0.837 | **0.871** | 0.642 | 0.751 |
| Phosphatases | 0.762 | **0.834** | 0.652 | 0.503 | 0.580 | **0.696** | 0.426 | 0.253 | 0.744 | **0.814** | 0.680 | 0.463 |
| Proteases | 0.562 | **0.738** | 0.667 | 0.597 | 0.316 | **0.545** | 0.445 | 0.357 | 0.483 | **0.700** | 0.641 | 0.590 |
| Protein kinases | 0.652 | **0.755** | 0.609 | 0.451 | 0.425 | **0.569** | 0.371 | 0.204 | 0.675 | **0.724** | 0.610 | 0.496 |
| Transporters | **0.354** | 0.237 | 0.082 | −0.018 | **0.125** | 0.056 | 0.007 | 0.000 | **0.220** | 0.190 | 0.032 | −0.030 |
| Mean | 0.604 | **0.605** | 0.460 | 0.403 | 0.391 | **0.396** | 0.246 | 0.196 | **0.571** | 0.566 | 0.451 | 0.403 |

RMSD analysis was performed across all 14 protein families and assessed all 6 engines. Following completion of the crystal-structure retrieval, every docked structure with an available co-crystal reference was evaluated, yielding 11,024 pose–crystal pairs across 2,064 structures (compared with the 1,053-structure subset available previously). Figure S3 shows the empirical cumulative distribution function (ECDF) of RMSD values per engine, where the y-axis represents the fraction of poses at or below a given RMSD threshold, with vertical lines marking the 1 Å and 2 Å thresholds. Our findings show RevDock with 73.3% ≤ 2.0 Å, a mean of 1.60 Å, and a median of 1.01 Å (Figure S3A). PandaDock, rerun with expanded sampling, was evaluated with 72.1% ≤ 2.0 Å, a mean of 2.22 Å, and the lowest median RMSD of any engine at 0.96 Å (Figure S3B). DiffDock was assessed with 56.4% ≤ 2.0 Å and a median of 1.50 Å (Figure S3C). AutoDock-GPU obtained 52.2% ≤ 2.0 Å and a median of 1.87 Å (Figure S3D). Boltz2 obtained 45.3% ≤ 2.0 Å and a median of 2.49 Å (Figure S3E). rDock was assessed with 40.7% ≤ 2.0 Å and a median of 2.50 Å (Figure S3F).

Protein family difficulty was assessed by pooling RMSD results across all engines per family. On the complete structure set we found that Transporters (66.8% ≤ 2.0 Å, N = 470) (Figure S4N), Ion channels (64.6%, N = 711) (Figure S4E), Molecular chaperones (62.4%, N = 956) (Figure S4G), and Nuclear receptors (62.0%, N = 1,016) (Figure S4H) performed the best across all 14 protein families. The protein families that did not perform well on this docking power assessment are Oxidoreductases (43.7%, N = 872) (Figure S4I), GPCRs (45.9%, N = 307) (Figure S4C), and Epigenetic enzymes (46.0%, N = 967) (Figure S4B). Glycosidases, previously excluded for insufficient coverage (N = 12), reached N = 1,054 pose–crystal pairs on the complete set and performed above the median at 61.0% ≤ 2.0 Å (Figure S4D). Notably, this ordering does not reproduce the pattern obtained on the earlier 1,053-structure subset, in which the membrane-embedded families ranked among the poorest performers; on complete data Transporters and Ion channels rank first and second, while GPCRs remain below median.

RevDock was assessed with 83.1% ≤ 2.0 Å in PPI targets, 82.8% for Molecular chaperones, 82.4% for Nuclear receptors, 77.6% for GPCRs, and 77.5% for Protein kinases; with performance dropping to 64.2% for Glycosidases, 64.5% for Proteases, and 58.6% for Ion channels (Figure 3). PandaDock, following the expanded-sampling rerun, reproduced poses reliably for PPI targets (87.3%), Molecular chaperones (84.0%), Transporters (84.0%), Ion channels (83.9%), and Nuclear receptors (82.9%), with weakest performance on Oxidoreductases (51.0%) and Epigenetic enzymes (59.5%) (Figure 3). DiffDock performance was competitive on Molecular chaperones (66.2%), Metalloenzymes (65.7%), and Protein kinases (64.6%), with performance weakest on GPCRs (30.0%) and Oxidoreductases (31.5%) (Figure 3). AutoDock-GPU performed best on Ion channels (77.6%) and Nuclear receptors (74.8%), with moderate performance between 25–70% across the other families (Figure 3). Boltz2 reached 73.4% on Ion channels and 70.5% on Metalloenzymes, declining to 20.9% for PPI targets and 23.2% for GPCRs (Figure 3). rDock reached 64.0% on Transporters and 60.2% on Nuclear receptors, declining to 26.5% for Proteases and 29.1% for Oxidoreductases (Figure 3).

**Figure 3:**
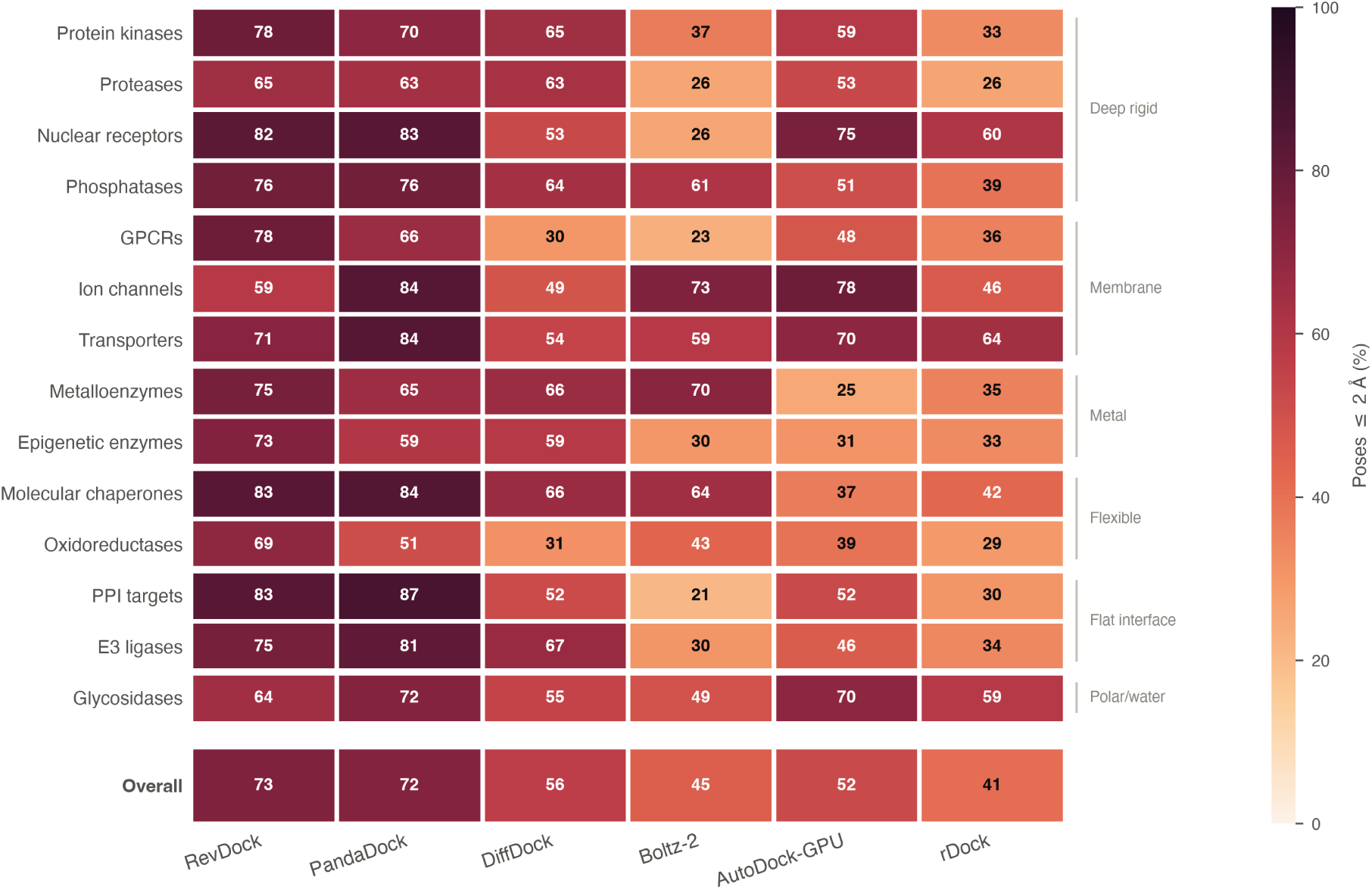
Docking power by engine and target family: percentage of predicted poses within 2 Å of the crystal ligand, over 11,024 pose–crystal pairs from 2,064 structures. Right-hand brackets group families by mechanistic class.

**Table 2:** Docking pose accuracy across all 14 protein families for six docking engines, computed on the complete set of locally available crystal structures (6,368 ligand SDFs and 6,368 protein PDBs). Median RMSD (Å) and percentage of poses with RMSD ≤ 2 Å are reported per protein family. Bold values indicate the best-performing engine per family per metric. A lower median RMSD and a higher % poses ≤ 2 Å indicate better pose reproduction.

| Protein Family | Median RMSD (Å) | | | | | | % Poses $\leq 2$ Å | | | | | |
| --- | --- | --- | --- | --- | --- | --- | --- | --- | --- | --- | --- | --- |
|  | RevDock | DiffDock | PandaDock | AutoDock-GPU | rDock | Boltz2 | RevDock | DiffDock | PandaDock | AutoDock-GPU | rDock | Boltz2 |
| E3 ligases | 0.86 | 1.20 | **0.79** | 2.17 | 3.31 | 9.16 | 75.0 | 66.7 | **81.5** | 45.8 | 33.9 | 30.4 |
| Epigenetic enzymes | **0.84** | 1.21 | 1.24 | 3.48 | 2.99 | 5.52 | **72.6** | 58.8 | 59.5 | 30.8 | 33.2 | 29.8 |
| GPCRs | **1.07** | 3.79 | 1.07 | 2.29 | 2.74 | 13.51 | **77.6** | 30.0 | 66.0 | 47.8 | 35.7 | 23.2 |
| Glycosidases | 1.52 | 1.62 | **0.84** | 1.28 | 1.63 | 2.05 | 64.2 | 54.5 | **72.2** | 69.6 | 58.8 | 49.0 |
| Ion channels | 1.90 | 3.06 | **0.60** | 0.74 | 2.18 | 0.97 | 58.6 | 48.7 | **83.9** | 77.6 | 46.2 | 73.4 |
| Metalloenzymes | 1.17 | **1.08** | 1.45 | 3.16 | 2.64 | 1.37 | **75.2** | 65.7 | 65.4 | 25.4 | 35.4 | 70.5 |
| Molecular chaperones | 0.83 | 1.13 | **0.60** | 2.73 | 2.35 | 1.44 | 82.8 | 66.2 | **84.0** | 37.0 | 42.3 | 64.3 |
| Nuclear receptors | **0.59** | 1.71 | 0.61 | 0.98 | 1.67 | 4.47 | 82.4 | 52.6 | **82.9** | 74.8 | 60.2 | 25.8 |
| Oxidoreductases | **0.94** | 7.12 | 1.89 | 2.80 | 3.20 | 2.51 | **69.3** | 31.5 | 51.0 | 39.4 | 29.1 | 43.3 |
| PPI targets | 1.03 | 1.88 | **0.78** | 1.80 | 2.96 | 9.04 | 83.1 | 51.9 | **87.3** | 52.2 | 30.2 | 20.9 |
| Phosphatases | **0.67** | 1.27 | 0.81 | 1.92 | 2.36 | 1.31 | **76.5** | 63.6 | 76.2 | 50.8 | 39.3 | 60.7 |
| Proteases | **1.15** | 1.45 | 1.30 | 1.93 | 3.25 | 4.28 | **64.5** | 62.7 | 63.5 | 52.8 | 26.5 | 25.9 |
| Protein kinases | **1.06** | 1.16 | 1.16 | 1.61 | 3.14 | 3.78 | **77.5** | 64.6 | 69.7 | 58.8 | 33.0 | 36.8 |
| Transporters | 0.81 | 1.80 | **0.63** | 1.12 | 1.55 | 1.46 | 70.8 | 53.9 | **84.0** | 69.9 | 64.0 | 58.5 |
| **Overall** | 1.01 | 1.50 | **0.96** | 1.87 | 2.50 | 2.49 | **73.3** | 56.4 | 72.1 | 52.2 | 40.7 | 45.3 |

Physical validity assessment was then conducted across all 11,036 poses generated from each engine and protein family. AutoDock-GPU was assessed with a pass rate of 98.1% (N = 1,649), rDock at 94.9% (N = 2,062), Boltz2 at 94.8% (N = 2,050), DiffDock at 94.0% (N = 1,803), PandaDock at 93.1% (N = 1,860), and RevDock at 73.2% (N = 1,605) (Figure 4). Seven poses timed out at the five-second limit and were excluded from the denominators. Five of the six engines therefore cluster within five percentage points of one another, and RevDock is the single clear outlier. RevDock’s failures are concentrated in Ion channels (22%), Glycosidases (57%), PPI targets (58%), E3 ligases (59%), Proteases (61%) and Protein kinases (65%), while it performs well on Nuclear receptors (98%), Molecular chaperones (91%) and Metalloenzymes (88%) (Figure 4). The remaining five engines exceed 84% in every family; the lowest individual cells are Boltz2 on Proteases (84%) and PandaDock on Metalloenzymes (85%). Because these five engines differ from one another by less than the spread within any single family, physical validity does not discriminate between them on this benchmark and should not be read as a ranking axis.

**Figure 4:**
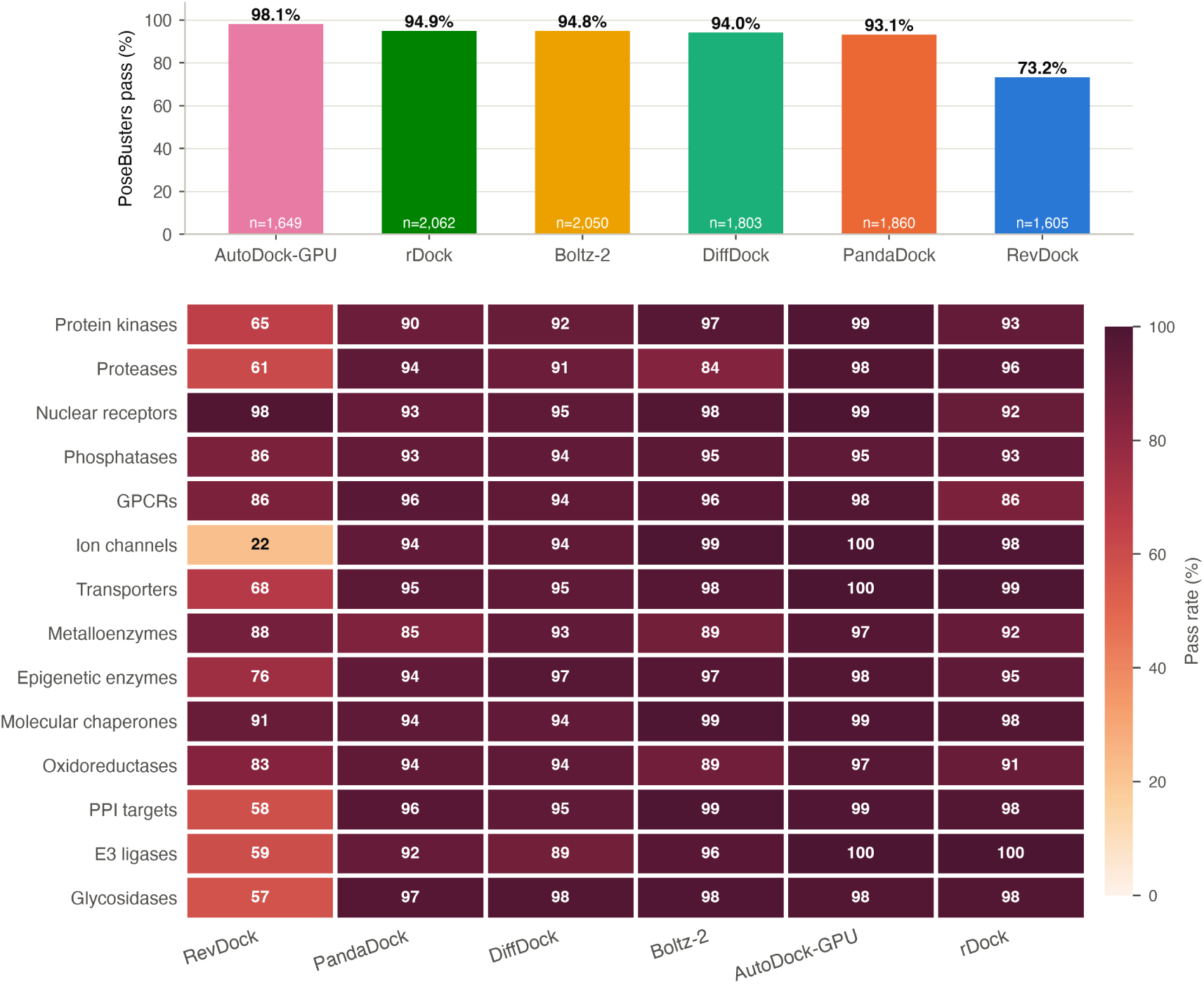
Physical validity by PoseBusters. Top: overall pass rate per engine, with the number of evaluable poses shown inside each bar. Bottom: pass rate by engine and target family. A pose passes only if every PoseBusters mol-mode check is satisfied; the seven poses of 11,036 that exceeded the five-second evaluation limit are excluded from the denominators.

**Figure 5:**
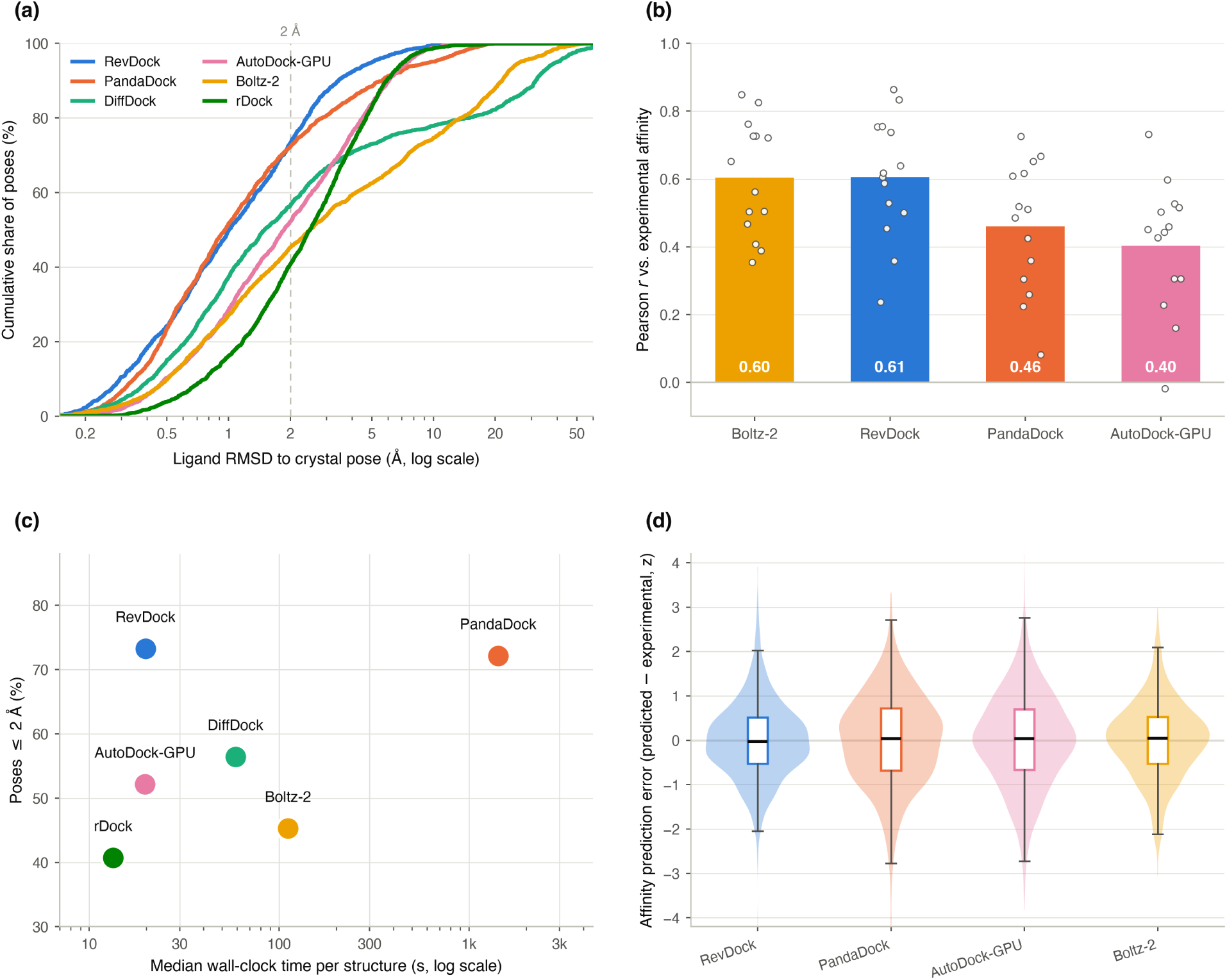
Consolidated performance summary across the four analytical axes. (a) Cumulative distribution of ligand RMSD for all six engines over the 11,024 pose–crystal pairs; the height of each curve at the 2 Å rule is the docking-power success rate of Table 2, and the shape away from that threshold shows how the engines differ in the near-native and grossly-wrong regimes. (b) Pearson *r* against experimental affinity: bars are the mean over families, open circles the 14 individual family values, so between-family spread is visible alongside the mean. (c) Cost against pose accuracy: median wall-clock time per structure versus percentage of poses within 2 Å. (d) Signed affinity prediction error, z-scored within family, as violin, box and median; centring away from zero indicates systematic bias and width indicates variance. Panels (b) and (d) are restricted to the four engines whose outputs are calibrated to an experimental affinity scale.

**Figure 6:**
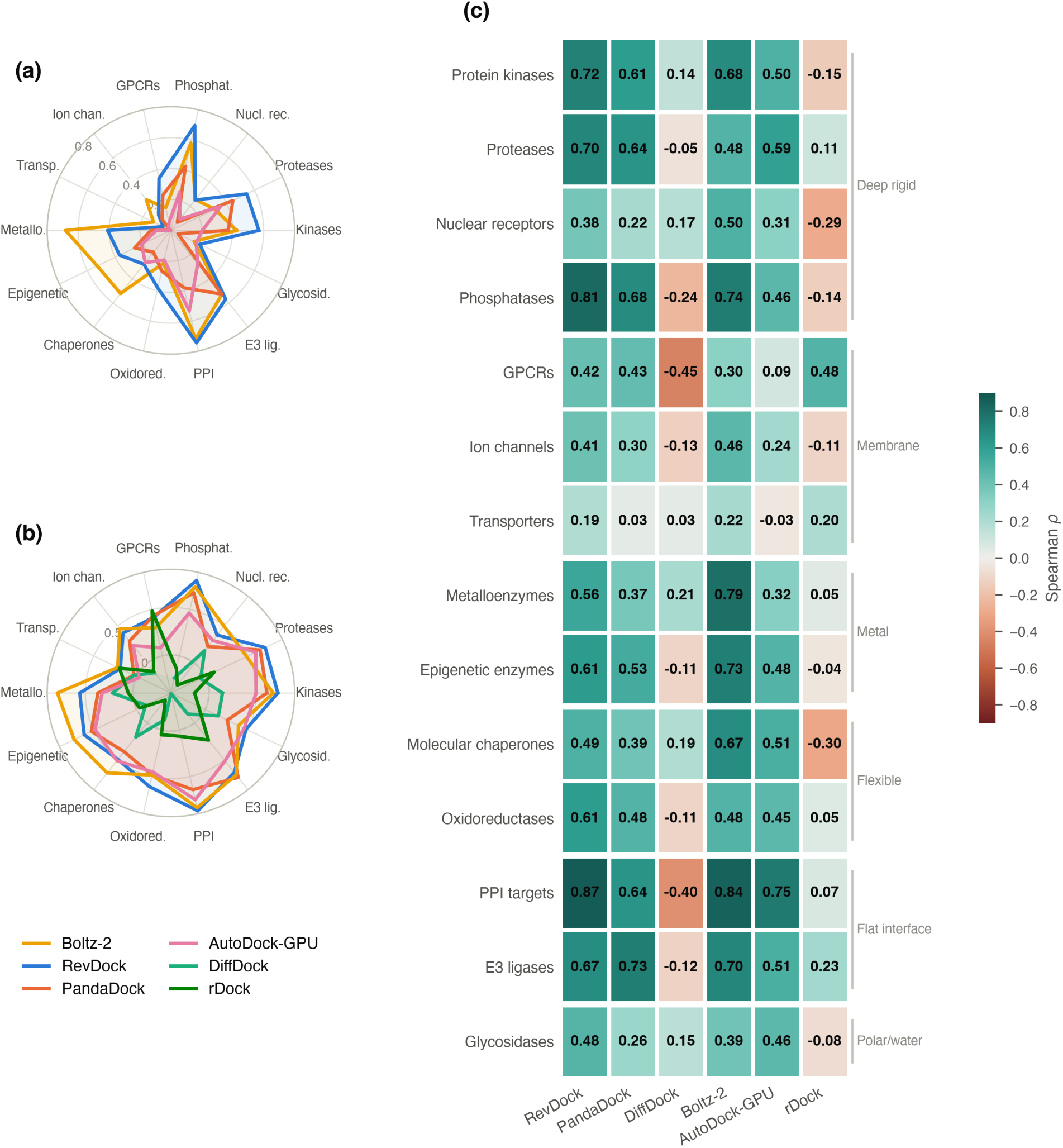
Family-level performance profiles on the two affinity axes. (a) Scoring power (*R*^2^) across the 14 target families for the four engines whose outputs are calibrated to an experimental affinity scale. (b) Ranking power (Spearman *ρ*) across the same families for all six engines, including the two Tier-4 rank-only engines. In both radars the informative quantity is the *shape* of each profile rather than its enclosed area: Boltz2 and RevDock reach almost identical means (Table 1) while peaking on different families, which is the central observation of this work. (c) The same ranking power as a signed matrix, so values and direction can be read directly; families in which an engine orders compounds inversely to experiment appear in red. Right-hand brackets group families by mechanistic class.

## 4 Discussion

Six engines were benchmarked across 14 protein families using four analytical axes. RevDock led pose accuracy overall with 73.3% ≤ 2.0 Å and a median of 1.01 Å, with PandaDock statistically indistinguishable at 72.1% and the lowest median RMSD of any engine (0.96 Å). Boltz2 and RevDock were statistically equivalent on affinity prediction, both with a mean r of approximately 0.6 and 12 of 14 families significant. PandaDock and AutoDock-GPU formed a second tier on affinity (mean r = 0.460 and 0.403 respectively), both significant in 12 of 14 families. On physical validity, five of the six engines cluster between 93.1% and 98.1% and are not meaningfully separable; RevDock is the sole outlier at 73.2%, its failures concentrated in Ion channels (22%) and the flat-interface families. Physical validity therefore functions as a floor that most engines clear rather than as a discriminating axis. Contrary to the pattern seen on the earlier partial-coverage subset, membrane-embedded families were not uniformly the hardest targets once every available co-crystal structure was evaluated: Transporters and Ion channels rank first and second on pooled pose accuracy, while GPCRs remain below median. Overall, no engine led on more than one axis, and the two engines with a learned component led on affinity while the two with the largest search budgets led on pose reproduction.

According to^13^, the greatest challenge to molecular docking is the limitations of the scoring function, which are a result of trading accuracy for speed with the use of approximations. Specifically, the force fields utilized in these scoring functions are classical approximations of the quantum mechanical description of molecular interactions, where solving this quantum mechanical description at the highest fidelity proves to be infeasible at a large scale^14^. Solving this molecular docking scoring accuracy limitation has somewhat been addressed with simplification of the quantum mechanics formulations; however, it introduces four key caveats in terms of scoring across different protein families and ligands of differing chemical backgrounds. These four limitations include, but are not limited to, the following: an assumption of rigid receptors, a pairwise additivity limitation, a limitation with fixed point charges, and an implicit solvent limitation. Assuming rigidity of the receptor removed the need to model the protein across multiple conformations. The Born-Oppenheimer approximation had been introduced as a way to model the quantum mechanics of a protein-ligand complex at scale, where instead of accounting for the motion of each electron, we can treat the electrons as an average cloud, where the quantum aspect of nuclear motion can be ignored^15^. In Lemkul 2020, he addresses this pairwise additive limitation as well as fixed charge limitation as a computational bottleneck where nonbonded interactions are the most expensive part of an MD simulation. The solution to this bottleneck is a pairwise additive setup with each atom in the pair being assigned a fixed charge^16^. The use of an implicit solvent model was explained in Izadi et al. 2015, where they claimed that the use of this model, as the average dielectric properties of water significantly reduced the cost of simulation in comparison to a discrete water model previously utilized^17^. What these four limitations have in common is that they are all solutions to solve for the computational bottleneck in regards to Quantum Mechanical modeling of protein-ligand binding.

According to the four limitations we have identified, we have classified all 14 protein families into six mechanistic groups based on which simplification is most severely stressed. The groups include Membrane Embedded Proteins, Metal Coordination Proteins, Deep Rigid Pocket Proteins, Flexible and Shallow Pocket Proteins, Large Flat Interface Proteins, and Polar Water-Dependent Proteins (see Table S2, Supporting Information).

AutoDock-GPU is a physics-based engine with no learned mitigation layer^18^. It utilizes a Lamarckian genetic algorithm to provide a global conformation search capable of escaping the local minima limitation observed with gradient descent^18^. The scoring function is depicted in the equation below:

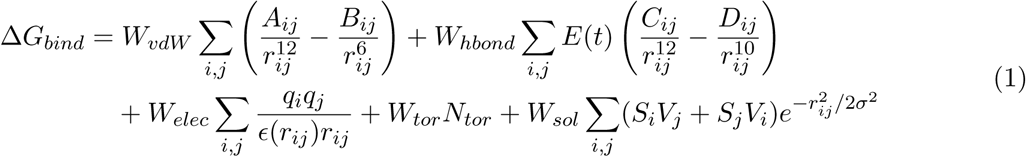

The weights depicted in this equation are fit by regression against a small training set of approximately 200 protein-ligand complexes, substantially smaller and less diverse than datasets such as PDBbind v2020. The weights are not case-specific for protein families; therefore, the scoring function will only reflect the protein families well represented by the training set. Autodock-GPU’s scoring function, as depicted above, is limited by our four assumptions that we have identified earlier: rigid receptor conformation, implicit solvent assumption, fixed point charge assumption, and pairwise additivity assumption. Affinity correlation analysis shows both scoring power assessment and rank ordering assessment, both with very large variances. Pearson r ranged from –0.02 with Transporters to 0.73 with PPI targets. Spearman *ρ* ranged from –0.03 with transporters to 0.75 with PPI targets. When comparing PPIs to the transporters, we notice a few things. PPI‘s are soluble proteins, unlike Transporters. The PPIs consist of relatively rigid hot spot geometry in contrast to transporters with an alternating access mechanism where the assumption of receptor rigidity becomes invalid. Lastly, although both PPIs and Transporters both do not consist of metal ions, the membrane environment in Transporters amplifies polarization effects that fixed charges in this engine miss. For pose reproduction assessment on the complete structure set, Ion channels performed the best with 77.6% ≤ 2.0 Å, followed by Nuclear receptors (74.8%) and Glycosidases (69.6%), while Metalloenzymes performed the worst with 25.4% ≤ 2.0 Å, followed by Epigenetic enzymes (30.8%). The Metalloenzyme result is the clearest single failure attributable to a named simplification: AutoDock-GPU treats Zn^2+^ as a fixed point charge, so the tetrahedral coordination geometry that positions the warhead carries no directional term in the scoring function, and the search is free to place chemically reasonable poses in the wrong orientation within the coordination shell. Notably, the deep-rigid-pocket families that this engine should suit best (Protein kinases 58.8%, Proteases 52.8%, Phosphatases 50.8%) sit mid-pack rather than at the top, indicating that pocket rigidity alone does not predict AutoDock-GPU’s pose accuracy. Physical validity assessment shows the highest pass rate of any engine at 98.1%, with every family at or above 95% and four at 99–100%. This reflects the Lamarckian genetic algorithm’s local-search step, which relaxes each candidate into a clean local minimum before scoring; AutoDock-GPU’s weakness on this benchmark is where it places the ligand, not the geometry of what it places.

Similar to AutoDock-GPU, rDock is also a physics-based engine with no learned mitigation layer^19^. Unlike the other docking engines, it utilizes a cavity-based search with a 6 Å radius compared to the docking grid box format, which is more efficient for sampling tight, well-defined pockets, but will fail for large flat interfaces. The scoring function is depicted with the following equation:

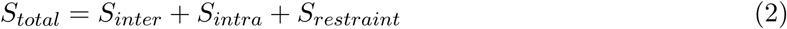

where

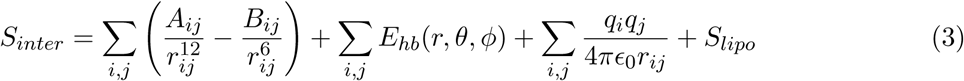

The equation captures the intermolecular score between the ligand and protein, the intramolecular ligand penalizing term that penalizes torsional strain and internal clashes to ensure ideal ligand conformational states, and an optional pharmacophoric or distance restraint. The equation effectively captures hydrophobic burial as modeled by the Gaussian lipophilicity term and ideal physical geometry with the intramolecular strain term. Fast sampling within a small cavity, specifically a ligand-centred 6 Å cavity, makes sampling efficient for well-defined pockets. Unlike RevDock and Boltz2, whose outputs are explicitly calibrated to experimental affinity scales, rDock’s dimensionless scoring function is not trained on binding affinity data. As a result *R*^2^ and Pearson r scoring power assessment could not be conducted for rDock. Spearman *ρ* rank ordering evaluation was performed as an exploratory analysis, which returned a mean *ρ* of 0.006 across the 14 families and reached significance in 4; no family showed a rank signal strong enough to support prospective ranking, consistent with a scoring function never fitted to affinity data. RMSD analysis on the complete structure set shows rDock with the lowest docking power of the six engines (40.7% ≤ 2.0 Å, median 2.50 Å), and an ECDF that rises later but no less steeply than the physics-based engines (Figure S3F): the distribution is not uniform, as the earlier partial-coverage analysis suggested, but shifted. rDock therefore does discriminate binding modes, simply less sharply than engines with either a learned rescoring layer or a larger search budget. At the protein-family resolution, Transporters performed the best with 64.0% ≤ 2.0 Å and Nuclear receptors second at 60.2%, while Proteases performed the worst at 26.5%, followed by Oxidoreductases (29.1%) and PPI targets (30.2%). The strong Transporter and Nuclear receptor results are consistent with rDock’s cavity-based search: both present a single, well-enclosed, largely hydrophobic cavity that a ligand-centred 6 Å sphere circumscribes accurately, and the Gaussian lipophilicity term has a clear gradient to follow. The PPI result is the converse case and the one the method predicts: a large flat interface is clipped by that same 6 Å cavity, so much of the true binding epitope lies outside the sampled volume. The poor Protease performance is not predicted by the mechanistic framework, which classifies proteases as deep and rigid; extended substrate-mimetic ligands spanning several subsites are plausibly the cause, since the S1–S4 groove is long rather than compact and is therefore also poorly circumscribed by a spherical cavity. In comparison to the other engines, rDock performed sixth of six on pose accuracy with an overall mean RMSD of 3.08 Å. Physical validity assessment shows rDock with an overall PoseBusters pass rate of 94.9%, and on the complete pose set no family falls below 86%; the previously reported PPI-target failure does not reproduce, and PPI targets now pass at 98%. rDock’s cavity restriction therefore costs it pose accuracy on flat interfaces without producing chemically invalid geometry.

PandaDock is a physics-based engine that utilizes a gradient descent algorithm to explore different ligand conformational states^20^; however, this technique is often unable to capture the global minima as there is no escape mechanism like the Lamarckian Genetic algorithm used in AutoDock-GPU or the Monte Carlo sampling methodology in rDock, and often gets stuck with a local minimum. Pandadock’s scoring function can be depicted in the equation below:

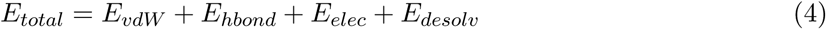

The scoring function makes all four assumptions that we have identified earlier (i.e., rigid receptor, pairwise additivity limitation, fixed point charge limitation, and implicit solvent assumption), meaning we will anticipate this to only perform well on deep rigid receptors, with no metal or strongly polar interactions, and independent interactions. We anticipate shallow pockets will not have enough conformation sampling time to be able to overcome the local minima. Under default settings this expectation was met and then some: affinity correlation was near-zero or negative across all 14 families (mean *r* = −0.001) and only 1.3% of poses fell within 2.0 Å. Rerunning the identical scoring function with an expanded search budget (v4.1.1; exhaustiveness 32, 10 retained poses, median 1,425 s per structure against ∼20 s for the other physics-based engines) changes both results completely: mean *r* rises to 0.460 with 12 of 14 families significant, and 72.1% of poses fall within 2.0 Å at a median RMSD of 0.96 Å, the lowest of any engine (Figure S8). Every family improves on both axes without exception.

This has a direct bearing on how the four simplifications should be read. Because the scoring function, the receptor preparation, the box definition, and the affinity ground truth are all identical between the two runs, none of the four approximations can account for the difference; only the adequacy of the conformational search changed. The default-parameter result was therefore not evidence about PandaDock’s scoring function at all. It also revises the earlier interpretation of the engine’s physical-validity behaviour. Under default sampling PandaDock returned the highest pass rate of any engine (96.1%) alongside the lowest pose accuracy, which indicated that local gradient-descent minimisation was reliably producing clean geometry in the wrong location. With expanded sampling the pass rate falls slightly to 93.1%, its lowest family being Metalloenzymes (85%), while pose accuracy rises to 72.1%. The engine now places ligands in genuinely occupied pockets, where steric constraints make ideal geometry harder to achieve; a small loss of physical validity in exchange for a fifty-fold gain in pose accuracy is the expected signature of that shift, and confirms that the original high pass rate was a symptom of unconstrained minimisation rather than a strength. At the family level PandaDock now performs best on PPI targets (87.3% ≤ 2.0 Å), Molecular chaperones (84.0%), Transporters (84.0%) and Ion channels (83.9%), and weakest on Oxidoreductases (51.0%) and Epigenetic enzymes (59.5%). The strong Molecular chaperone result is the most informative of these, since the flexible HSP90 lid was the specific case predicted to defeat a gradient-descent search; with sufficient restarts the engine locates the correct lid-adjacent pose, indicating that the barrier was the search landscape rather than the rigid-receptor treatment of the lid itself.

RevDock, our GNINA-derived engine, utilizes both physics-based scoring and applies a CNN rescoring layer that is trained on the CrossDocked2020 set, where it applies a CNN rescoring layer that transforms the protein-ligand complex into a 3D voxelized grid of atom density maps, from which CNN affinity is predicted in pKd units^6^. It has two key measures of affinity, VINA-based physics score and CNN affinity score. For this study, we will use CNN affinity. Since RevDock utilizes Vina style scoring, all four limitations of the quantum mechanical simplification remain applicable at the pose generation level. With the CNN scoring based on the CrossDocked2020 set, we anticipate limitations across protein families to be associated with representation in the training set. The CNN rescoring layer likely mitigates some of the limitations we have identified, particularly the pairwise additivity limitation, as the CNN learns from thousands of crystal structures where cooperative interactions are encoded and fixed point charges, since the CNN will implicitly learn the electrostatic patterns from the crystal structures.

Affinity correlation analysis shows RevDock with a very strong scoring power assessment and ranking power assessment. RevDock performed very well with the best scoring protein families, PPI targets, and Phosphatases, with a Pearson r above 0.80. The poor performance of Transporters (r = 0.24) and Ion channels (r = 0.36) out of the 14 protein families can be well explained by the fact that PPI targets and Phosphates are well represented in PDBbind-v2016. Both proteins are characteristic of their soluble nature as well as deep, well-defined pockets. Although PPI targets are characteristic of a hydrophobic burial pattern that the pairwise additivity cannot properly capture, and Phosphatases consist of a charged active site. The well-represented nature of both protein families in the CrossDocked2020 set allows the model to learn about interconnected interactions in PPI targets as well as the electrostatic patterns in Phosphatases. For Transporters and Ion Channels, their failure mode can largely be characterized by the fact that they are membrane-embedded proteins, which are both not well represented in the CrossDocked2020 set and also not able to be empirically captured due to the implicit solvent assumption. RMSD analysis also shows strong pose reproduction with Phosphatases with 94% of poses ≤ 2.0 Å. It is worth noting that E3 ligases reached 100% but with a sample size of only 6, a definitive conclusion cannot be made with this family. Reflecting the affinity correlation analysis, Ion channels did not perform well with only 20% of poses ≤ 2.0 Å. Although we had strong results for both affinity correlation and RMSD analysis, the physical validity evaluation underperformed, with 73.2% of poses passing, the only engine of the six to fall below 93%. At the protein family resolution, pass rates range from 22% (Ion channels) at the low end to 98% (Nuclear receptors) at the high end, with Glycosidases (57%), PPI targets (58%), E3 ligases (59%) and Proteases (61%) also well below the benchmark-wide norm. The per-family physical validity assessment is based on the representation of each family in the CNN training set; both PPI targets and Ion Channels are underrepresented in the set compared to Nuclear receptors, which are well represented. This is because the CNN rescoring layer is the learned representation of conformational geometry.

DiffDock, unlike AutoDock-GPU and RevDock, is a pure learning-based engine, with no physics-based equation application. Diffdock specifically is a generative diffusion model that uses neural networks as its core computational components^8^. Trained on the PDBbind-v2020 set, Diffdock was trained with a forward propagation process that aimed to add noise to the co-crystal structures along three specific axes: translational noise, rotational noise, and torsional noise^8^. The backpropagation step aimed to train the model to denoise the structure back to its original co-crystal structure. The model then outputs a confidence score for each sampled pose. The model is effective in diverse pose sampling, blind pocket identification, and generalizable representations. Similar to the CNN rescoring layer in RevDock, the assessment of each protein family is dependent on the representation of the protein family in PDBbind-v2020. Therefore, flexible targets, insoluble proteins, and novel binding modes will not be well captured by this engine.

Similar to rDock, DiffDock’s scoring function is not calibrated to experimental affinity scales^8^. As a result, only ranking power was assessed as an exploratory analysis, calculating Spearman *ρ*. Results show PPI targets (*ρ* = –0.395, p = 0.001), Phosphatases (*ρ* = –0.237, p = 0.008), and Metalloenzymes (*ρ* = 0.214, p = 0.009) as the most significant protein families out of the 14 evaluated; yet no family showed consistent affinity signal. RMSD results show protein kinases performing the best with 90% of all poses ≤ 2.0 Å. On the other end, Transporters performed the worst with 6% of poses ≤ 2.0 Å. The strong performance of protein kinases can be explained by the fact that it is the most heavily represented family in PDBbind v2020. Blind pocket identification used in DiffDock applies well in this case since the ATP cleft sits at the interface of the N and C-terminal lobes, making it clearly accessible from the protein surface. Meanwhile, Transporters are sparsely represented in the training set. The binding site is deeply embedded in the membrane, where Diffdock’s diffusion process has no mechanism to identify the pocket inside the transmembrane bundle. Physical validity assessment shows DiffDock with a strong PoseBusters pass rate of 94.0%. All 14 protein families sit above the 80% threshold, with E3 ligases the lowest at 89%. This performance is characterized by the graph neural network ligand representation that inherently encodes chemical validity; training on crystal structures allows the model to be able to learn the distribution of physically valid poses from the training set.

Boltz2 is a pure model-based algorithm, which utilizes a three-step process for molecular docking. It takes a protein sequence and uses EvoFormer, which utilizes Multiple Sequence Alignment (MSA) to be able to build the protein representation of the sequence^7^. The second step is the cofolding process, where the 3D coordinates for both protein and ligand are iterated over using invariant point attention^7^. This process was trained with PDB co-crystal structures. The last step is affinity prediction, where a regression model is trained using experimental affinity data from ChEMBL and PDBbind^7^. The use of the protein sequence in the first step allows the rigid receptor assumption to be bypassed, although this step can be limited by protein sequences with shallow MSA, as these novel proteins would not be able to be properly mapped with Evoformer. The cofolding step and affinity prediction step will be able to bypass both the pairwise additivity limitation and the fixed point charge limitation, as the model trained for this step is able to learn both the codependent interactions as well as the electrostatic interactions of each protein system. These steps will largely be limited by the distribution of each protein family across the training step.

Affinity correlation results show Boltz2 performing fairly well, with PPI targets performing the best with a Pearson r of 0.85, *R*^2^ of 0.72, and Spearman *ρ* of 0.85. On the other end, Transporters performed the worst with Pearson r of 0.35, *R*^2^ of 0.12, and Spearman *ρ* of 0.22. The strong performance of PPI targets can be associated with the strong co-evolutionary signal that Evoformer is able to capture with the PPI hotspots, where Evoformer is effectively able to map the protein topology. Bypassing the rigid receptor assumption is a factor in the strong performance, as PPI inhibitors often induce a slight conformational change at the hot spot pocket upon binding. For Transporters, the implicit solvent assumption becomes fully exposed, as Boltz2 is not properly able to model environments of differing dielectric terms. Transporters are very sparsely represented in the training set. RMSD analysis was conducted on Boltz2 with the caveat that since docking was not conducted on the crystal structure, alignment was necessary to properly compute RMSD. Post-alignment analysis shows Proteases as the best-performing family with 86% of poses ≤ 2.0 Å with a sample size of 21. On the contrary, Molecular Chaperones performed the worst with 15% of poses ≤ 2.0 Å. The results with Proteases are in line with why the performance of affinity correlation was also strong. In the case of Molecular Chaperones, it does reveal that the flexible lid problem is not solved by co-folding. This is likely due to the model averaging across lid conformations during training rather than taking into account that each conformation needs to be accounted for the full nature of the protein. Different proteins in the Molecular Chaperone family (i.e., HSP90, HSP70, GRP78, etc.) all have different binding site geometries where Evoformer is unable to model due to the heterogeneity in this specific family. Physical validity assessment shows Boltz2 with a PoseBusters pass rate of 94.8%, placing it with the four physics-based engines rather than below them. Protein family-specific analysis shows every family above 84%, the lowest being Proteases (84%), Metalloenzymes (89%) and Oxidoreductases (89%). We note that this figure supersedes an earlier estimate of 88.0% obtained before ligand bond orders were restored from the input SMILES; the previously reported Phosphatase deficit (72%) was an artefact of that extraction step rather than a property of the predicted poses, and Phosphatases in fact pass at 95%. Co-folding therefore does not incur a physical-validity penalty relative to classical docking, and Boltz2’s weaknesses on this benchmark are confined to pose placement and affinity ranking on membrane-embedded families.

## 5 Conclusion

In this study, we benchmarked six docking and co-folding engines across 14 protein families along four analytical axes: scoring power, ranking power, docking power, and physical validity. Rather than treating each protein family as an independent benchmark, we classified all 14 families into six mechanistic groups based on which of four scoring-function simplifications, rigid receptor, pairwise additivity, fixed point charges, and implicit solvent, is most severely stressed by that family’s binding site character. This framework accounts for the majority of the performance differences observed across engines and protein families in this study, and shifts the question from “which engine is best” to “which structural or physical property of the target is this engine’s scoring function equipped to handle.”

Engines with a learned rescoring or generative component, RevDock’s CNN affinity layer and Boltz2’s sequence-based co-folding, consistently outperformed engines relying on unmodified physics-based scoring functions (AutoDock-GPU, rDock, PandaDock) on affinity correlation, while engines with robust global conformational search were more competitive on pose reproduction. No single engine performed best across all four axes and all 14 families; instead, performance tracked closely with the four structural and physical characteristics identified above, regardless of which specific family or engine was under consideration. Deep, rigid, soluble pockets with independent contacts and no metal coordination (e.g., protein kinases, proteases) were reproduced well by nearly every engine tested, while membrane-embedded families with cooperative, many-body binding character (GPCRs, ion channels, transporters) were poorly served by all six engines, reflecting the implicit-solvent and rigid-receptor assumptions shared across their scoring functions.

A methodological result follows from the PandaDock rerun. Under default sampling, PandaDock returned near-zero affinity correlation (mean *r* = −0.001) and 1.3% of poses ≤ 2.0 Å; with expanded conformational sampling and no change to its scoring function, the same engine reaches mean *r* = 0.460 and 72.1% of poses ≤ 2.0 Å. Because the scoring function is identical between the two runs, the difference is attributable entirely to search adequacy. This is a caution for benchmark studies generally: a result reported as a scoring-function limitation may instead reflect an under-converged conformational search, and the two are not separable without a sampling-convergence check.

For practitioners selecting a docking or co-folding engine for a specific target, these results suggest that the deciding factor is not an engine’s aggregate benchmark score, but whether that engine’s scoring function architecture addresses the specific structural or physical property, receptor flexibility, cooperative binding, electrostatic polarization, or membrane solvation, that dominates the target of interest. Extending this mechanistic framework to additional protein families, and to targets lacking high-resolution co-crystal structures, is a natural next step toward anticipating engine performance ahead of running a full benchmark.

## Supporting Information

Protein family characterization table summarizing structural hallmarks, cooperative binding character, electrostatic environment and solvent/membrane context for all 14 protein families; classification of the 14 families into six mechanistic groups by the dominant quantum-mechanical simplification stressed; per-engine affinity calibration scatter plots (Figure S1); per-family affinity calibration (Figure S2); per-engine RMSD cumulative distributions (Figure S3); per-family RMSD cumulative distributions (Figure S4); computational cost and run reliability (Figure S5); evaluable coverage per family by analytical axis (Figure S6); ranking power for every engine × family pair with confidence intervals (Figure S7); effect of conformational sampling on PandaDock (Figure S8); engine performance profiles across families (Figure S9); ranking power heatmap (Figure S10); performance profiles by method tier (Figure S11); distribution of prediction error by engine (Figure S12); pooled affinity calibration (Figure S13); scoring power heatmap for all six engines (Figure S14); and ranked ranking-power forest plot (Figure S15) (PDF).

## Acknowledgments

This research was funded internally by Revilico Inc. No external grant, funding agency, or compute-credit program supported this specific study.

