## Supplementary Information for "Structural Context Determines Docking Engine Performance: A Family-Stratified Benchmark of Six Engines"

### Supporting Information for: Structural Context Determines Docking Engine Performance: A Family-Stratified Benchmark of Six Engines

Kristoffer Alejo[1], Sarah Fisher[1], Tejaswan Kalluri[2], Balkrushna More[2], Hrishikesh Rajgure[2], Pritam Kumar Panda[3], Christopher Korban[1], and Christian Chung*[1]

[1]Revilico Inc., Los Angeles, California, USA
[2]Onelab Ventures Pvt. Ltd, Pune, Maharashtra, India
[3]Department of Anesthesiology, Perioperative and Pain Medicine, Stanford University School of Medicine, Stanford, California, USA

**Table S1:** Characterization of the 14 protein families across four docking-relevant properties corresponding to the four quantum mechanical simplifications. Families are ordered by mechanistic group.

| Protein Family | Conformational Dynamics (Rigid Receptor) | Cooperative Binding Character (Pairwise Additivity) | Electrostatic Environment (Fixed Point Charges) | Solvent/Membrane Context (Implicit Solvent) | Ref |
|---|---|---|---|---|---|
| **Protein Kinases** | DFG-in/out loop switching; glycine-rich P-loop collapses over inhibitor; mild induced fit at hinge region | Hydrophobic contacts and hinge H-bonds predominantly independent; pairwise approximation most valid of any family | No metal ions; moderate polar environment at hinge; fixed charges adequate for ATP-competitive chemotypes | Soluble; aqueous dielectric correct; modest water displacement from ATP pocket | [1;2] |
| **Proteases** | Rigid catalytic triad; minimal backbone movement upon binding; most conformationally stable family | Extended beta-strand H-bond pattern approximately additive; S1–S4 subsites contribute independently | No metal ions; active-site electrostatics modeled via fixed partial charges | Soluble; aqueous dielectric correct; active site solvation well-modeled by implicit solvent | [3–5] |
| **Nuclear Receptors** | Large hydrophobic LBD with rigid core; helix-12 repositions between agonist and antagonist states | Hydrophobic burial of polycyclic scaffold largely additive; mild cooperativity at coactivator interface | No metal ions; predominantly hydrophobic; fixed charges adequate for steroid-like scaffolds | Soluble; aqueous dielectric correct; large hydrophobic cavity with minimal ordered water network | [6] |

*Continued on next page*

Table S1 – *continued*

| **Protein Family** | **Conformational Dynamics (Rigid Receptor)** | **Cooperative Binding Character (Pairwise Additivity)** | **Electrostatic Environment (Fixed Point Charges)** | **Solvent/Membrane Context (Implicit Solvent)** | **Ref** |
|---|---|---|---|---|---|
| **Phosphatases** | WPD loop closes over active site upon substrate binding; allosteric sites show conformational change | Mild electrostatic cooperativity between catalytic Cys and Arg; pairwise approximation moderately stressed | Highly charged active site; catalytic Cys and Arg coordinate phosphate; fixed charges underestimate polarization | Soluble; aqueous dielectric correct; highly solvated active site with ordered water molecules partially displaced upon binding | [7;8] |
| **GPCRs** | Seven TM helices cycle between inactive and active states with up to 14 Å movement at TM6; ECL2 gates orthosteric pocket; allosteric sodium modulates conformation | Orthosteric contacts moderately cooperative; simultaneous TM helix interaction creates many-body character; allosteric sodium coordination inherently cooperative | Allosteric sodium (Asp2.50) creates charged TM environment; fixed charges miss sodium-dependent electrostatic modulation | Membrane-embedded; $\varepsilon \approx 2$–$4$ inside membrane vs. aqueous $\varepsilon \approx 80$; detergent crystallization misrepresents native environment | [9–11] |
| **Ion Channels** | Multimeric pore cycles through open, closed, and inactivated states; pore geometry completely different in each state; gating involves large coordinated helix movements | Pore blocking requires simultaneous interaction with four symmetrically arranged subunits; inherently many-body; pairwise terms cannot decompose | Selectivity filter contains highly charged residues coordinating ions; fixed charges miss ion-dependent electrostatic environment | Membrane-embedded; same dielectric mismatch as GPCRs; ion permeation driven by electrochemical gradient absent from all calculations | [12] |
| **Transporters** | Alternating access cycles through outward-facing, occluded, and inward-facing conformations with 10–20 Å helix movements; each state presents different binding site geometry | Substrate binding involves cooperative TM helix changes; thermodynamics depend on full conformational cycle, not a single state | Charged substrate binding residues in TM domain; membrane dielectric amplifies polarization effects that fixed charges cannot capture | Membrane-embedded; lipid bilayer $\varepsilon \approx 2$–$4$ fundamentally misrepresented by aqueous assumption | [13–16] |
| **Metalloenzymes** | Active site geometry adjusts slightly upon inhibitor binding; coordinating side chains reorient; mild induced fit at coordination shell | Zinc coordination involves four simultaneous groups (His-His-Glu/Asp-warhead) in tetrahedral arrangement; coordination energy cannot be decomposed into pairwise terms | Zinc is highly polarizable; electron density redistributes upon warhead coordination; fixed $Zn^{2+}$ charge severely underestimates coordination bond character | Soluble; aqueous dielectric correct; water displacement from coordination shell upon warhead binding not captured by implicit solvent | [17] |

*Continued on next page*

Table S1 – *continued*

| Protein Family | Conformational Dynamics (Rigid Receptor) | Cooperative Binding Character (Pairwise Additivity) | Electrostatic Environment (Fixed Point Charges) | Solvent/Membrane Context (Implicit Solvent) | Ref |
|---|---|---|---|---|---|
| **Epigenetic Enzymes** | Bromodomain pocket rigid and well-defined; HDAC active site has mild loop flexibility; HAT/HMT binding involves moderate conformational change | Bromodomain contacts predominantly additive; HDAC zinc coordination cooperativity identical to metalloenzymes; HAT/HMT has charged cofactor environment | Bromodomain: no metal, fixed charges adequate. HDAC: zinc-dependent, same polarization failure as metalloenzymes. HAT/HMT: charged cofactor environment | Soluble; aqueous dielectric correct; bromodomain acetyl-lysine pocket has ordered water network partially displaced upon binding | [18–22] |
| **Molecular Chaperones** | HSP90 flexible lid (residues 94–126) opens and closes over ATP binding site; different inhibitors stabilize different lid conformations; lid closure involves cooperative hydrophobic collapse | Lid closure is a multi-step dynamic process progressing from rapid local remodeling to slow concerted hydrophobic burial over the nucleotide pocket | No metal ions; mixed polar/hydrophobic character; fixed charges adequate for adenine-mimetic scaffolds but less accurate for polar lid-interacting groups | Soluble; aqueous dielectric correct; dynamic solvation state changes between open lid (solvated) and closed lid (buried) conformations | [23–25] |
| **Oxidoreductases** | Cofactor-dependent conformational changes alter pocket volume and geometry; substrate binding loop shows moderate flexibility; P450s have flexible substrate access channel | Cofactor-inhibitor-protein three-body interactions show mild cooperativity; competitive inhibitors interact simultaneously with both cofactor and protein residues | Cofactor (NAD(P)H, FAD, or heme) introduces charged/aromatic environment adjacent to inhibitor binding site; fixed charges miss cofactor-dependent polarization | Soluble; aqueous dielectric correct; cofactor presence or absence changes water network occupancy in a way implicit solvent cannot model | [26–28] |
| **PPI Targets** | Hot spot pockets show mild induced fit; MDM2-type pockets deepen slightly upon inhibitor binding; flat featureless PPIs have minimal conformational change | Interface binding inherently cooperative; hot spot residues contribute non-additively; burial of multiple aromatic residues cannot be decomposed into pairwise contacts; interface desolvation cooperative | Charged interface residues contribute to hot spot electrostatics; polarization effects at charged residues not captured by fixed charges | Soluble; aqueous dielectric correct; cooperative displacement of ordered interfacial water is a many-body solvation event not captured by pairwise implicit solvent | [29–31] |

*Continued on next page*

Table S1 – *continued*

| **Protein Family** | **Conformational Dynamics (Rigid Receptor)** | **Cooperative Binding Character (Pairwise Additivity)** | **Electrostatic Environment (Fixed Point Charges)** | **Solvent/Membrane Context (Implicit Solvent)** | **Ref** |
|---|---|---|---|---|---|
| **E3 Ligases** | Deep hydrophobic warhead pockets adjacent to protein recruitment interface; ternary complex formation involves induced fit not captured by rigid binary docking | PROTAC ternary complex formation involves simultaneous cooperative binding of three components; binary docking cannot model this; molecular glue binding also cooperative | No metal ions; hydrophobic warhead pockets (e.g. thalidomide pocket of CRBN); fixed charges adequate for small molecule warhead chemotypes | Soluble; aqueous dielectric correct; cooperative interface desolvation upon ternary complex formation not modeled by implicit solvent | [32–35] |
| **Glycosidases** | Relatively rigid active site; conserved catalytic dyad/triad position; mild distortion of sugar ring geometry upon substrate binding | Cooperative water-mediated H-bond networks dominate the polar active site; non-additive many-body polarization missed by independent pairwise force field terms | Positively charged iminosugar inhibitors pair with catalytic carboxylates (Asp/Glu); pH-dependent stabilization across ER and lysosomal environments | Soluble; aqueous dielectric correct; structural active-site water network treated as featureless continuum, omitting specific water displacement thermodynamics | [36–39] |

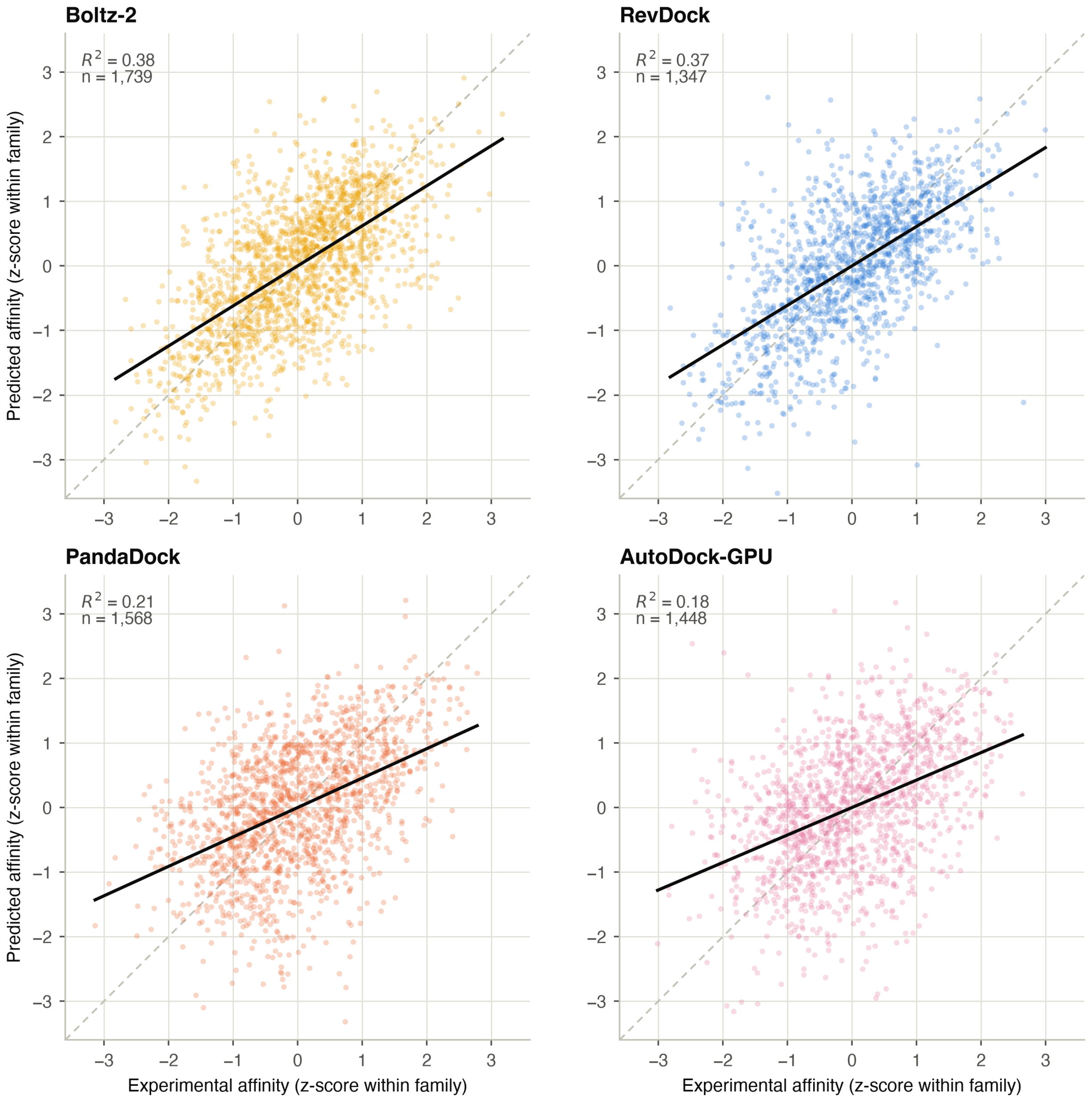


**Figure S1:** Predicted vs. experimental binding affinity, by engine. Panels: (A) Boltz2, (B) RevDock, (C) PandaDock, (D) AutoDock-GPU. Values are z-scored within each protein family so engines reporting on different native scales are directly comparable. Dashed line is the ideal predictor ($y = x$); solid line is the OLS fit.

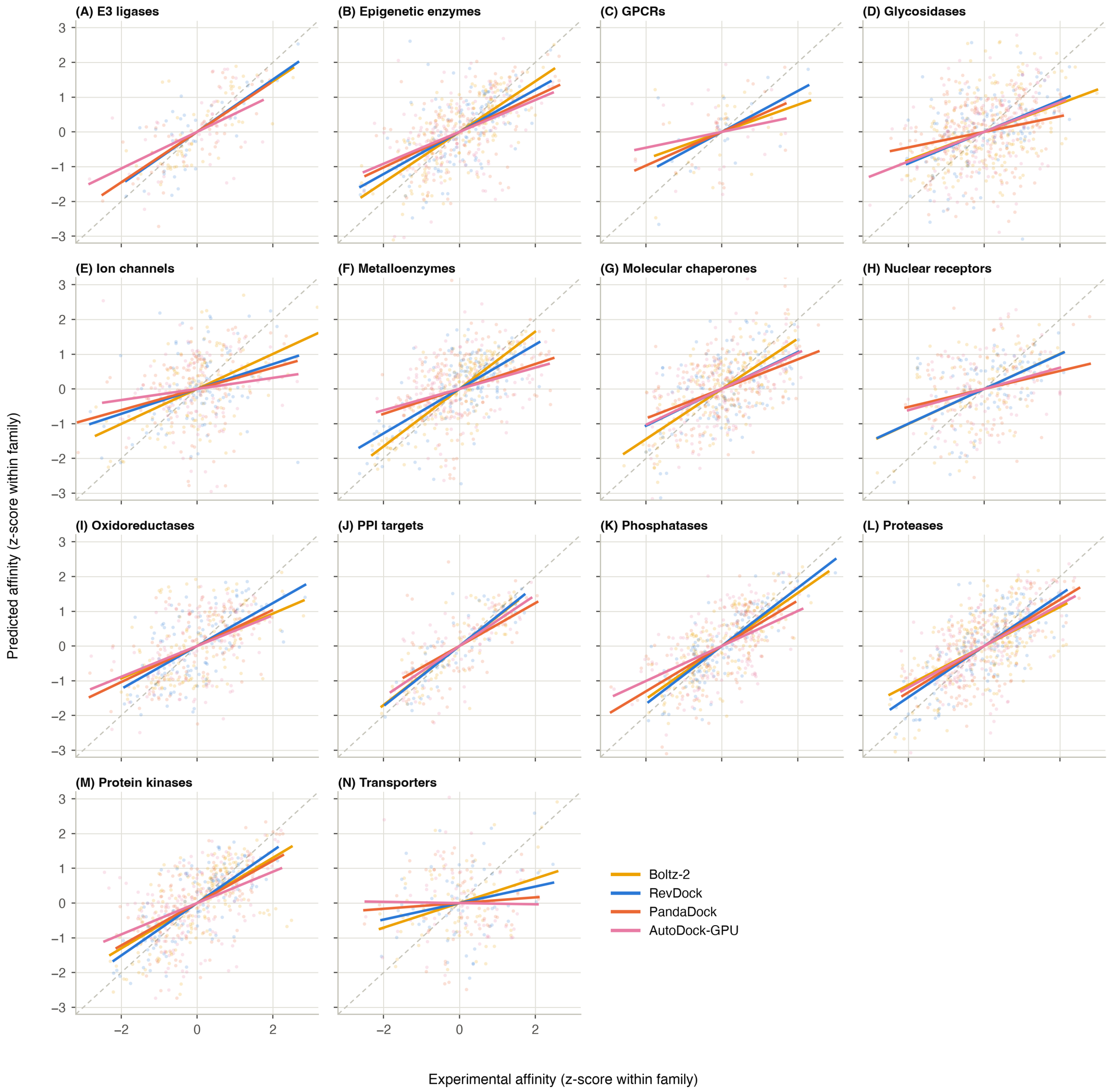


**Figure S2:** Affinity calibration by target family, panels (A)–(N) in alphabetical family order. Coloured lines are per-engine OLS fits; the dashed diagonal is the ideal predictor. Panels share axes, so slope differences are directly comparable across families.

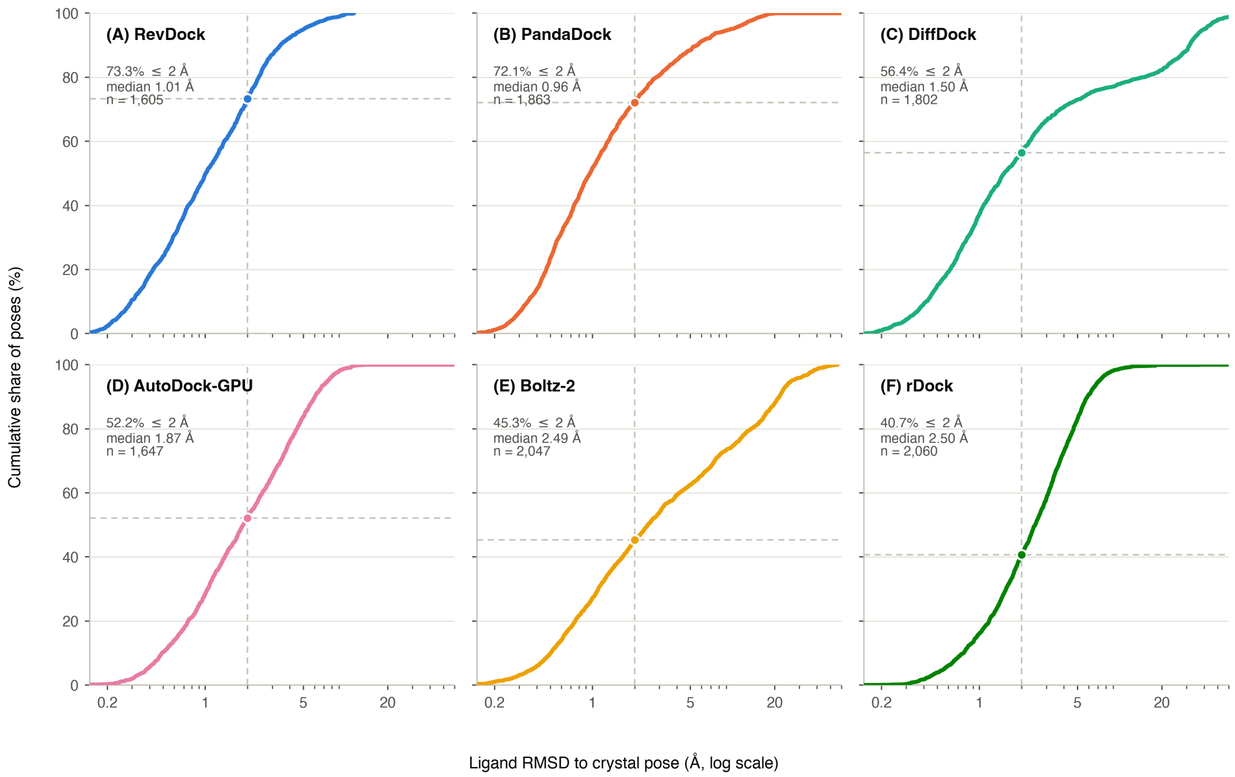


**Figure S3:** Empirical cumulative distribution of ligand RMSD, by engine: (A) RevDock, (B) PandaDock, (C) DiffDock, (D) AutoDock-GPU, (E) Boltz2, (F) rDock. Dashed lines mark the 2 Å docking-power threshold; the marker gives the docking-power success rate reported in the main text.

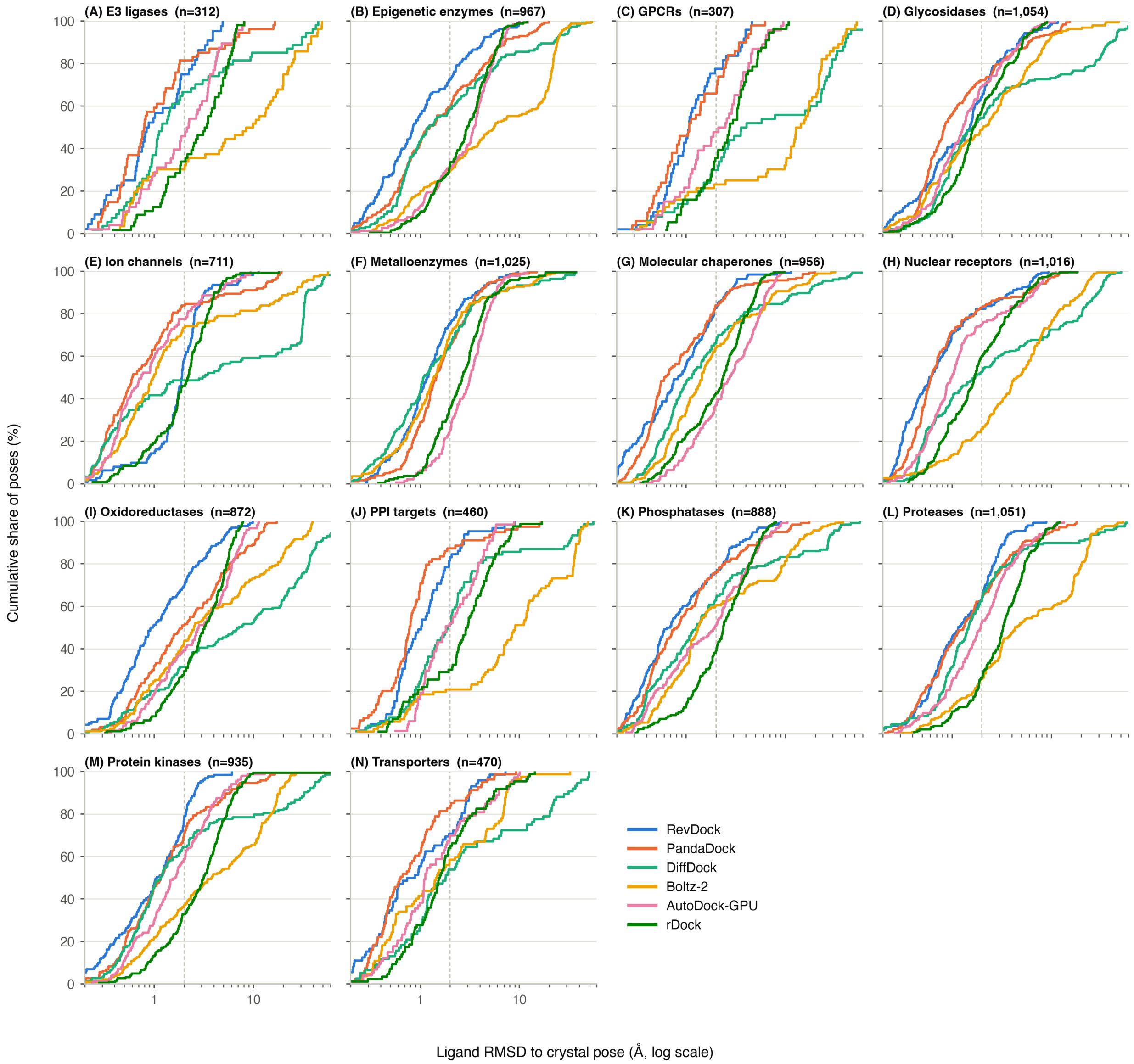


**Figure S4:** Empirical cumulative distribution of ligand RMSD by target family, panels (A)–(N). All six engines are overlaid within each panel; the dashed line marks the 2 Å threshold.

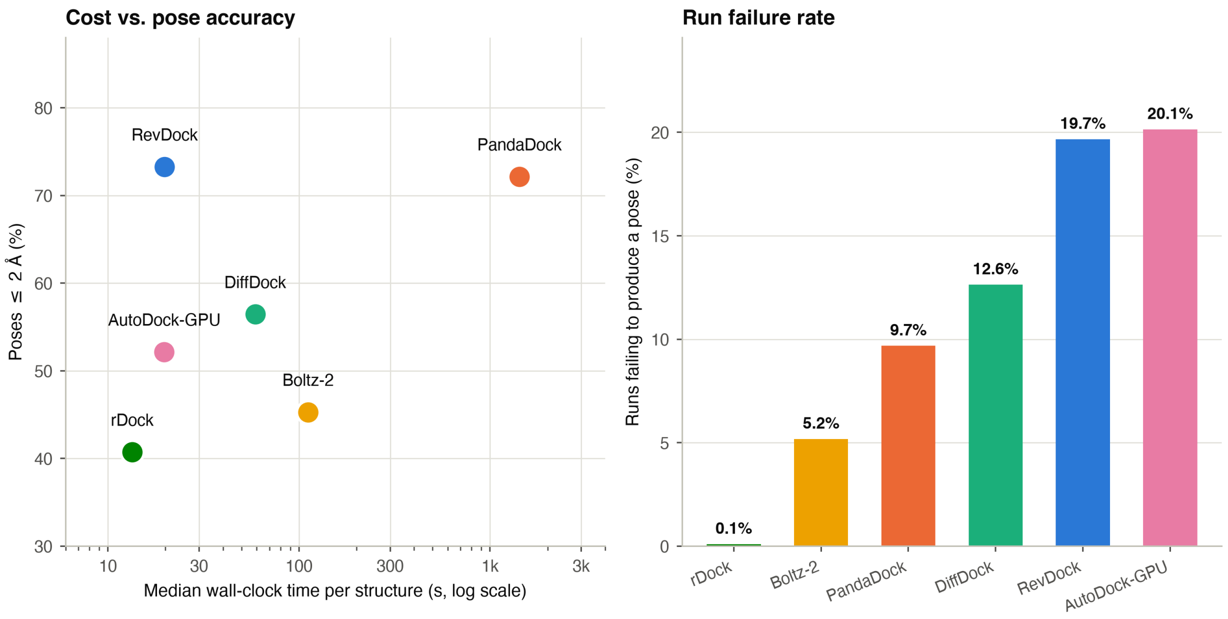


**Figure S5:** Computational cost and run reliability. Left: median wall-clock time per structure against docking-power success rate. Right: percentage of the 12,420 attempted runs that produced no usable output pose.

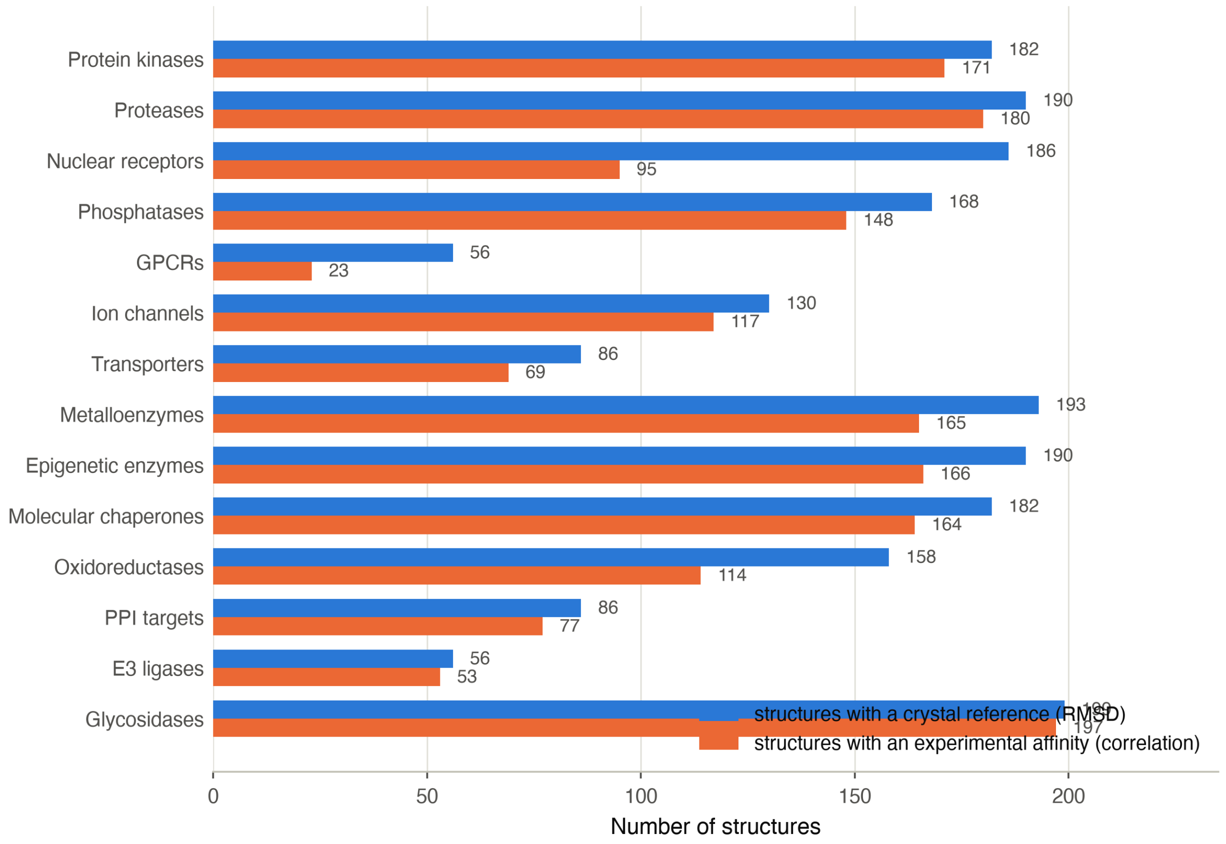


**Figure S6:** Evaluable coverage per target family, by analytical axis. Docking power and affinity correlation are powered on different subsets of the benchmark, so per-family sample size differs between the docking-power and affinity-correlation tables in the main text.

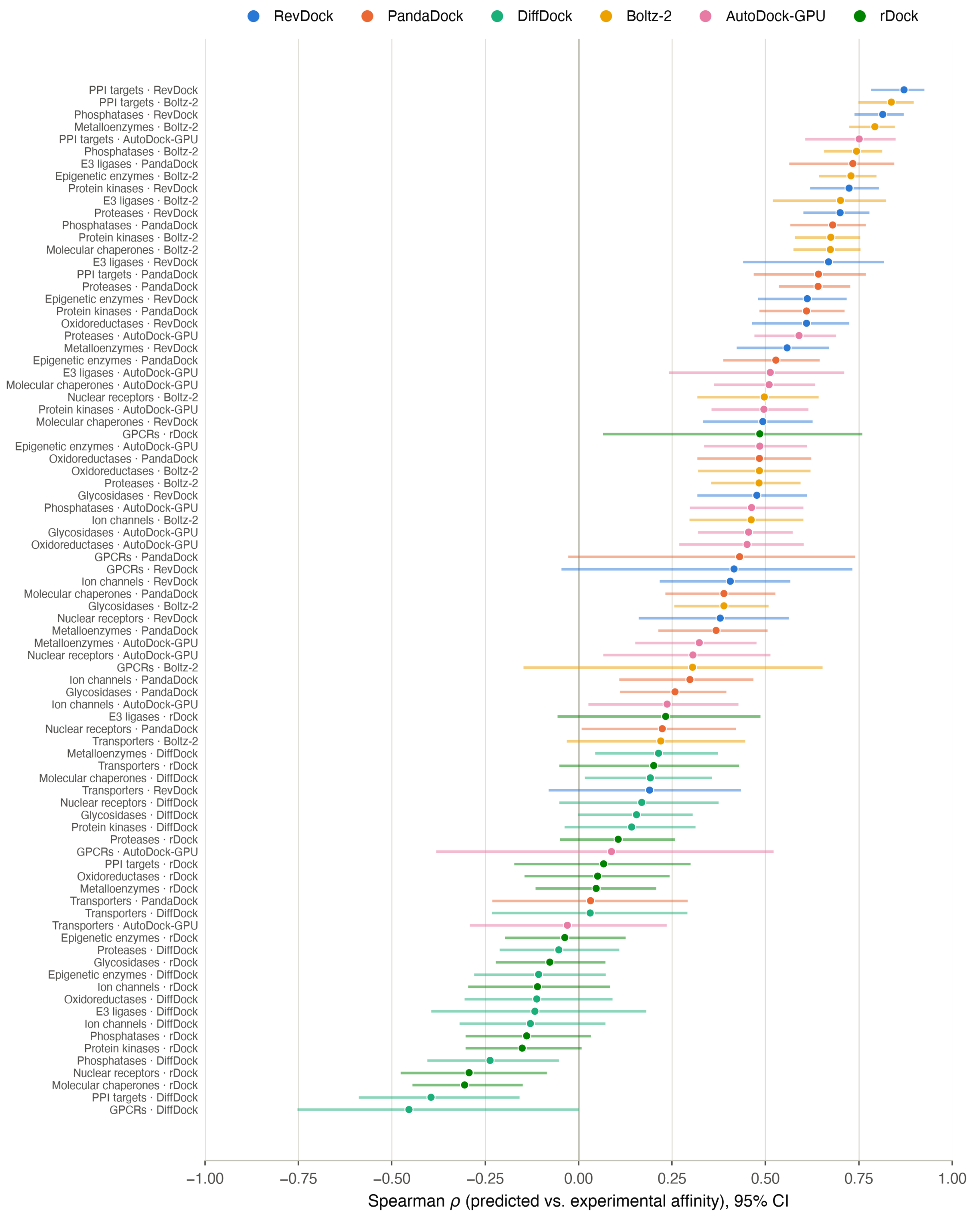


**Figure S7:** Ranking power for every engine × target-family pair, sorted by Spearman $\rho$ with 95% confidence intervals (Fisher $z$, SE $= 1.06/\sqrt{n-3}$). The vertical rule at $\rho = 0$ separates pairs with a positive rank signal from those without.

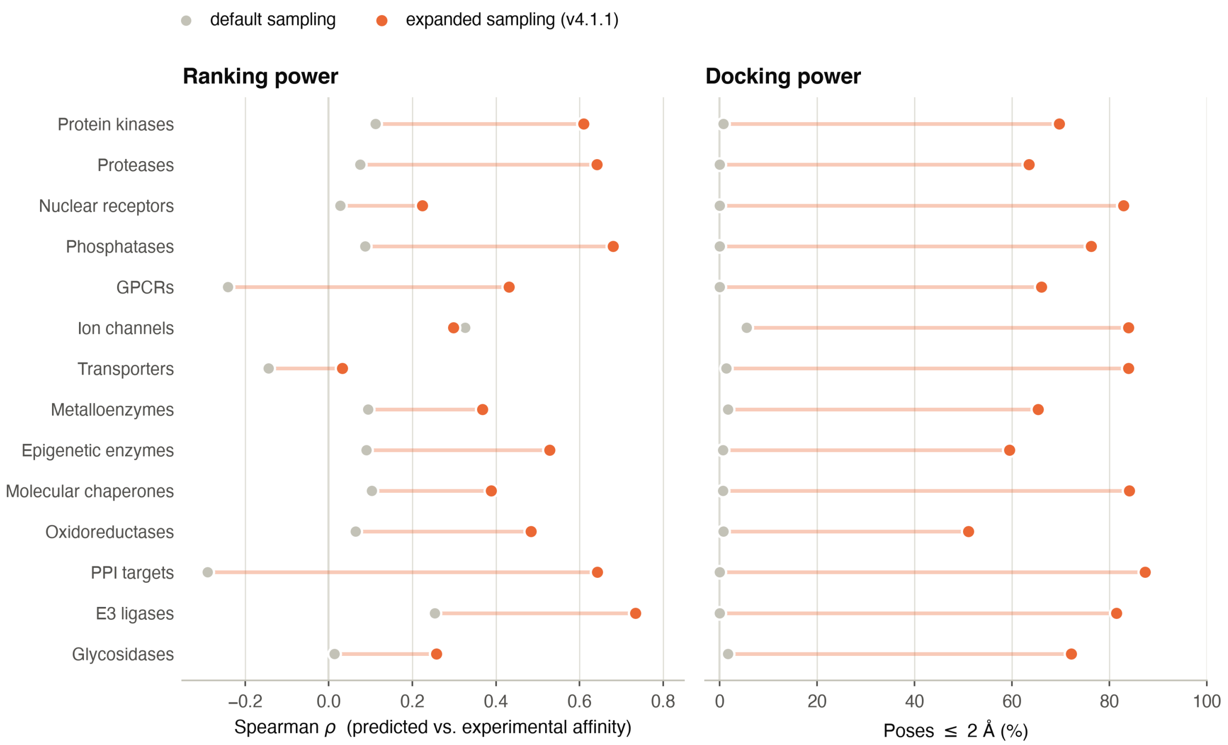


**Figure S8:** Effect of conformational sampling on PandaDock, holding the scoring function fixed. Grey points are the default-parameter run; coloured points are the expanded-sampling rerun (v4.1.1; exhaustiveness 32, 10 retained poses per run). Left: ranking power (Spearman $\rho$ against experimental affinity). Right: docking power (percentage of poses within 2 Å). Every family improves on both axes. Because the scoring function, receptor preparation, box definition and affinity ground truth are identical between the two runs, the difference is attributable to search adequacy alone.

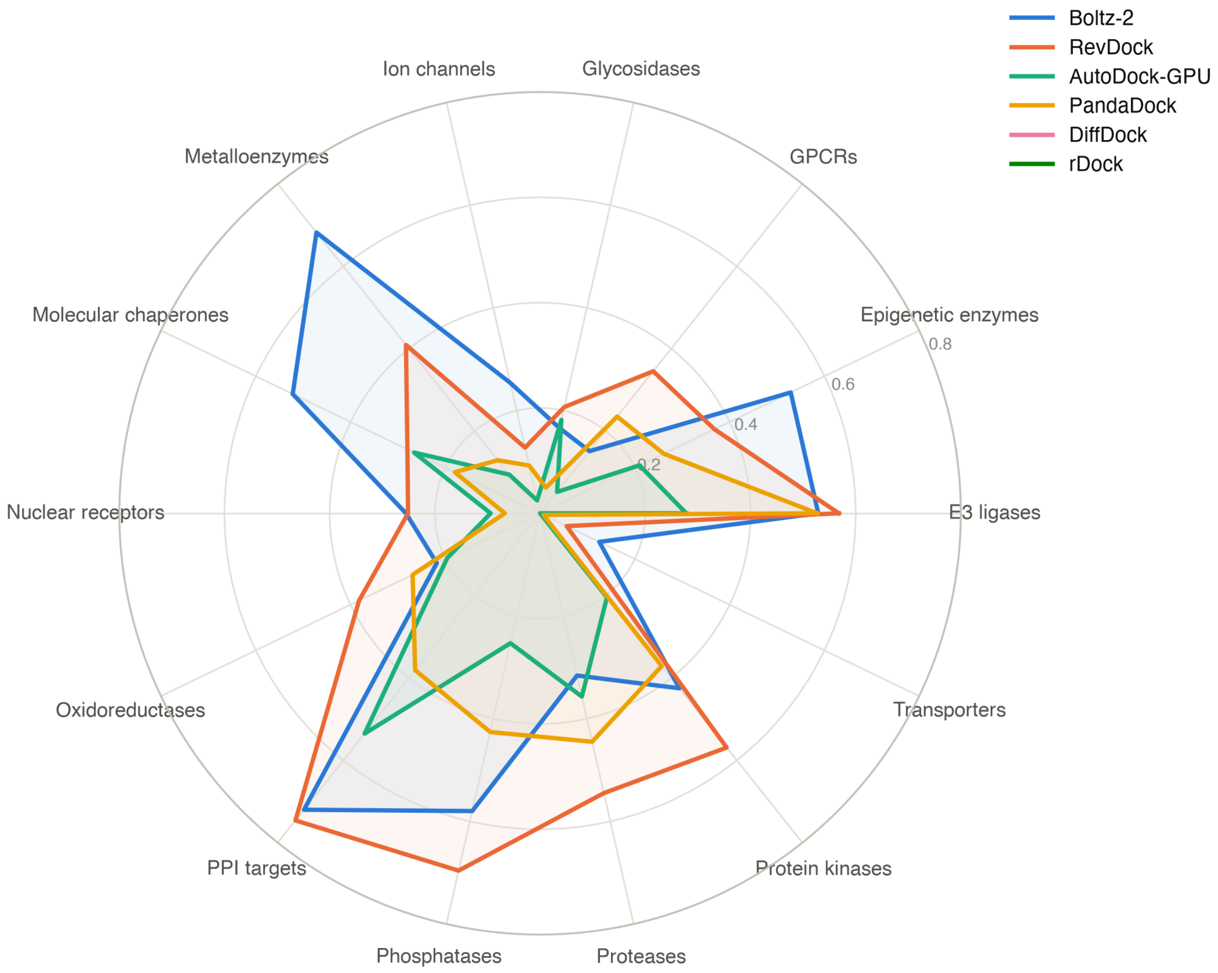


**Figure S9:** Engine performance profile across target families. Each spoke is one protein family and each line one engine, with radial distance the $R^2$ between predicted and experimental affinity. The shape of each profile, rather than its overall size, is what distinguishes the engines: Boltz2 and RevDock peak on different families despite near-identical mean $R^2$.

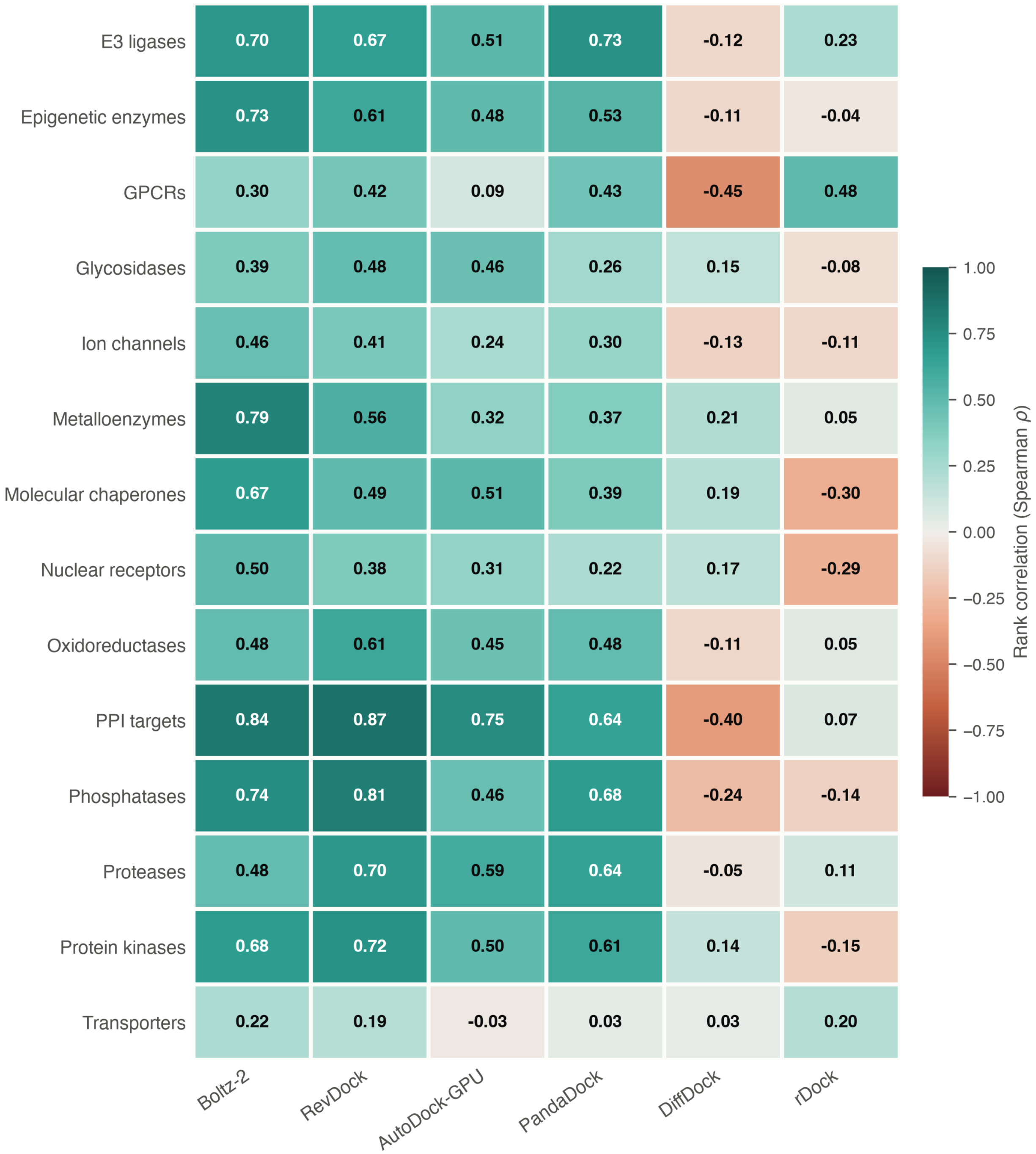


**Figure S10:** Ranking power by target family and engine, as Spearman $\rho$ between predicted and experimental affinity. Unlike the $R^2$ matrix in the main text, this scale is signed, so families where an engine ranks compounds in the wrong order are visible as negative cells. All six engines are shown, including the two Tier-4 rank-only engines.

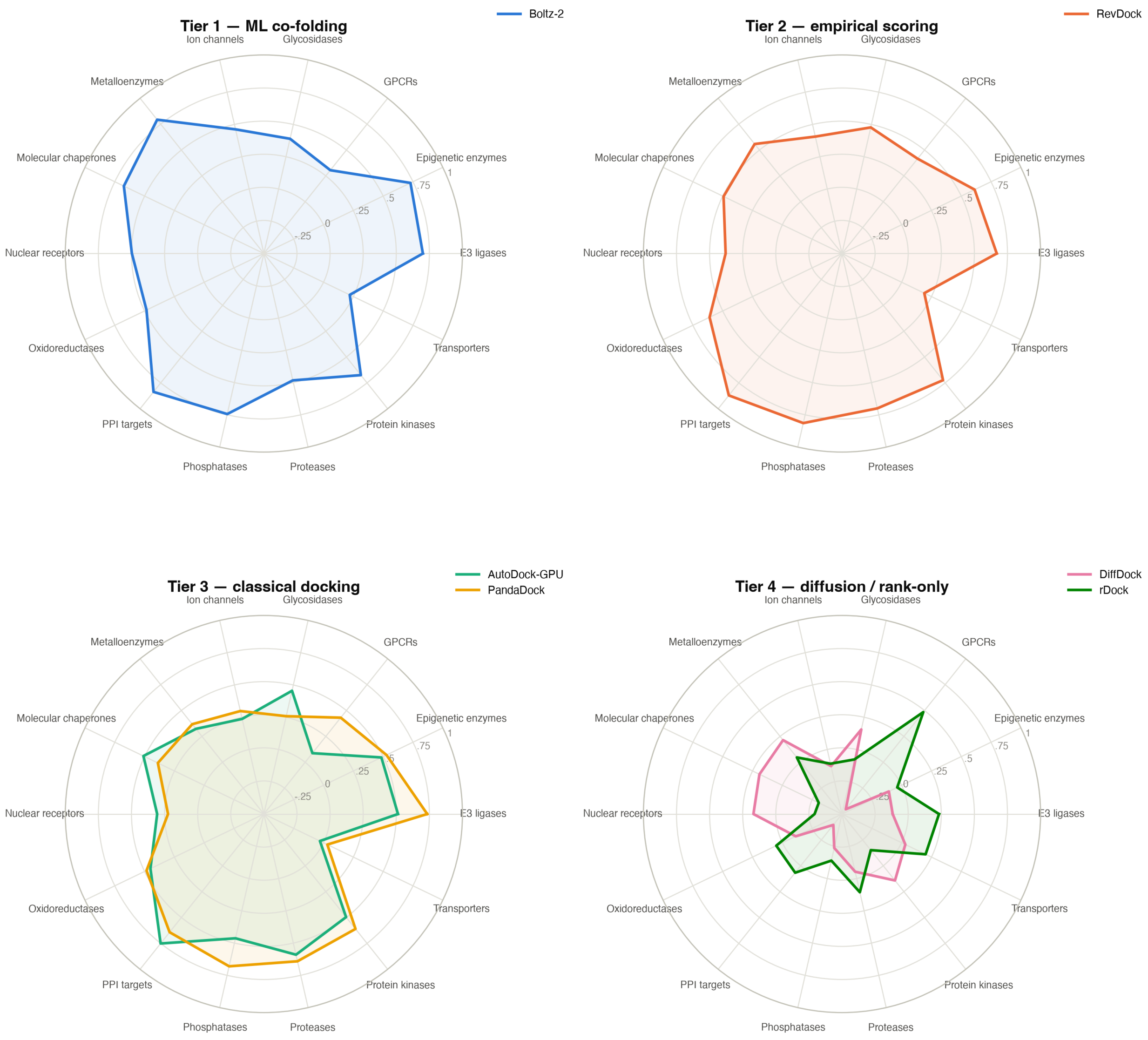


**Figure S11:** Ranking power by method tier. Panels separate the four method tiers defined in the main text so that engines are compared against others making the same class of prediction; radial distance is Spearman $\rho$ per family.

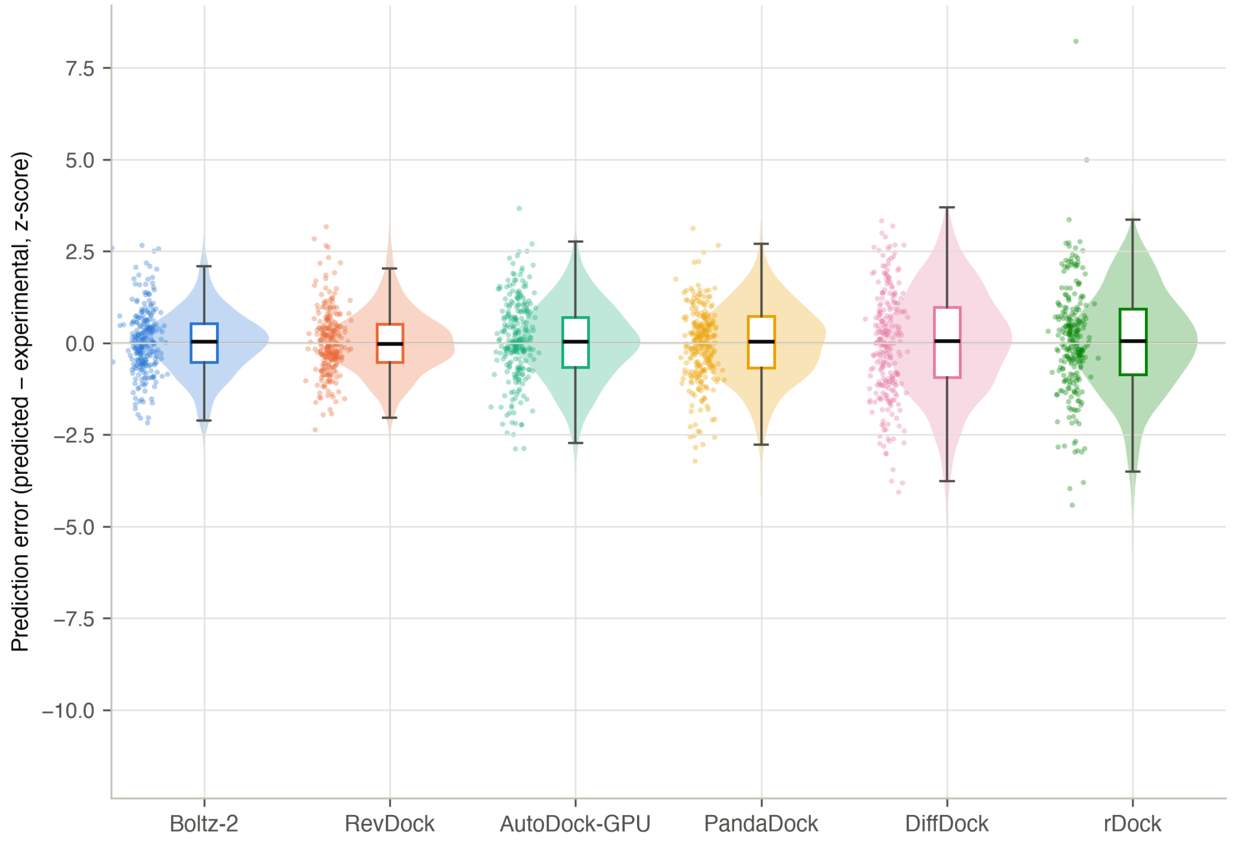


**Figure S12:** Distribution of signed prediction error (predicted − experimental, z-scored within family) per engine, shown as violin, box and jittered points. This separates bias from variance: an engine centred on zero with wide spread is noisy, whereas an offset centre indicates a systematic over- or under-prediction of affinity.

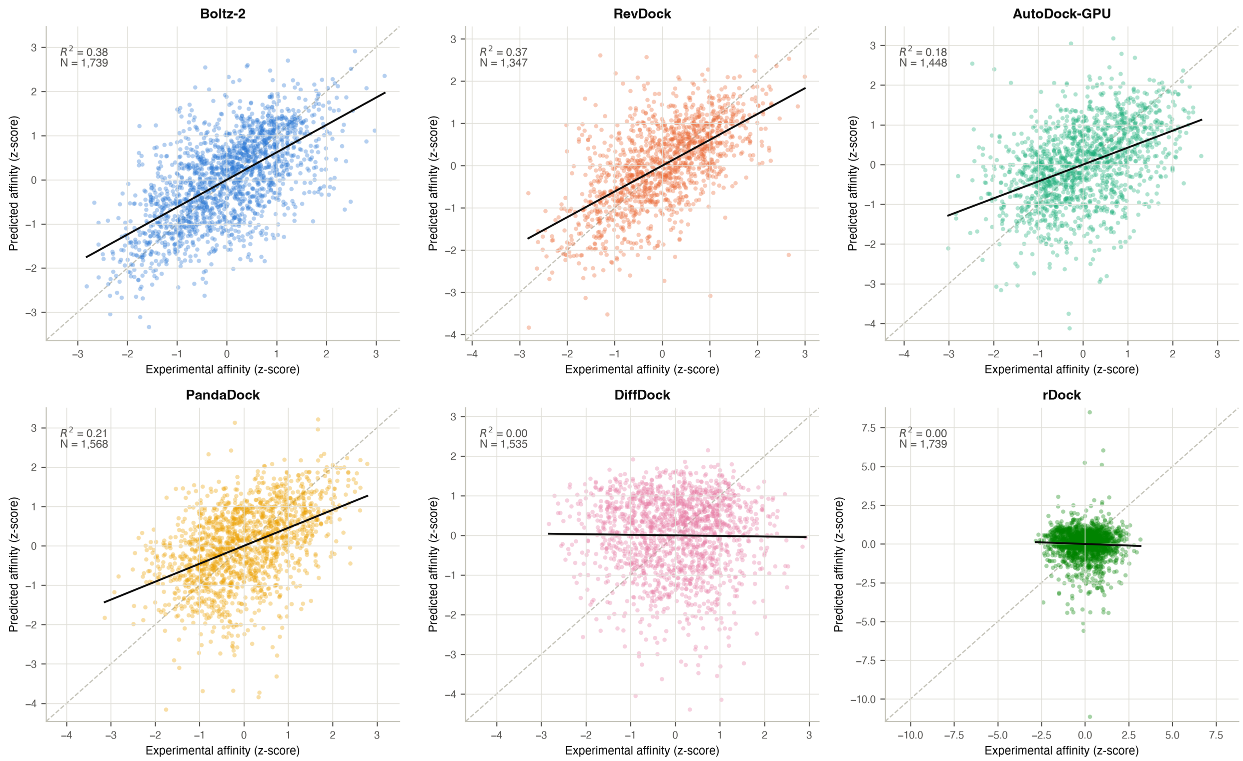


**Figure S13:** Predicted vs. experimental affinity per engine, pooled across all families, with an OLS fit and the $y = x$ ideal. This is the pooled counterpart to the per-family view in Figure S1.

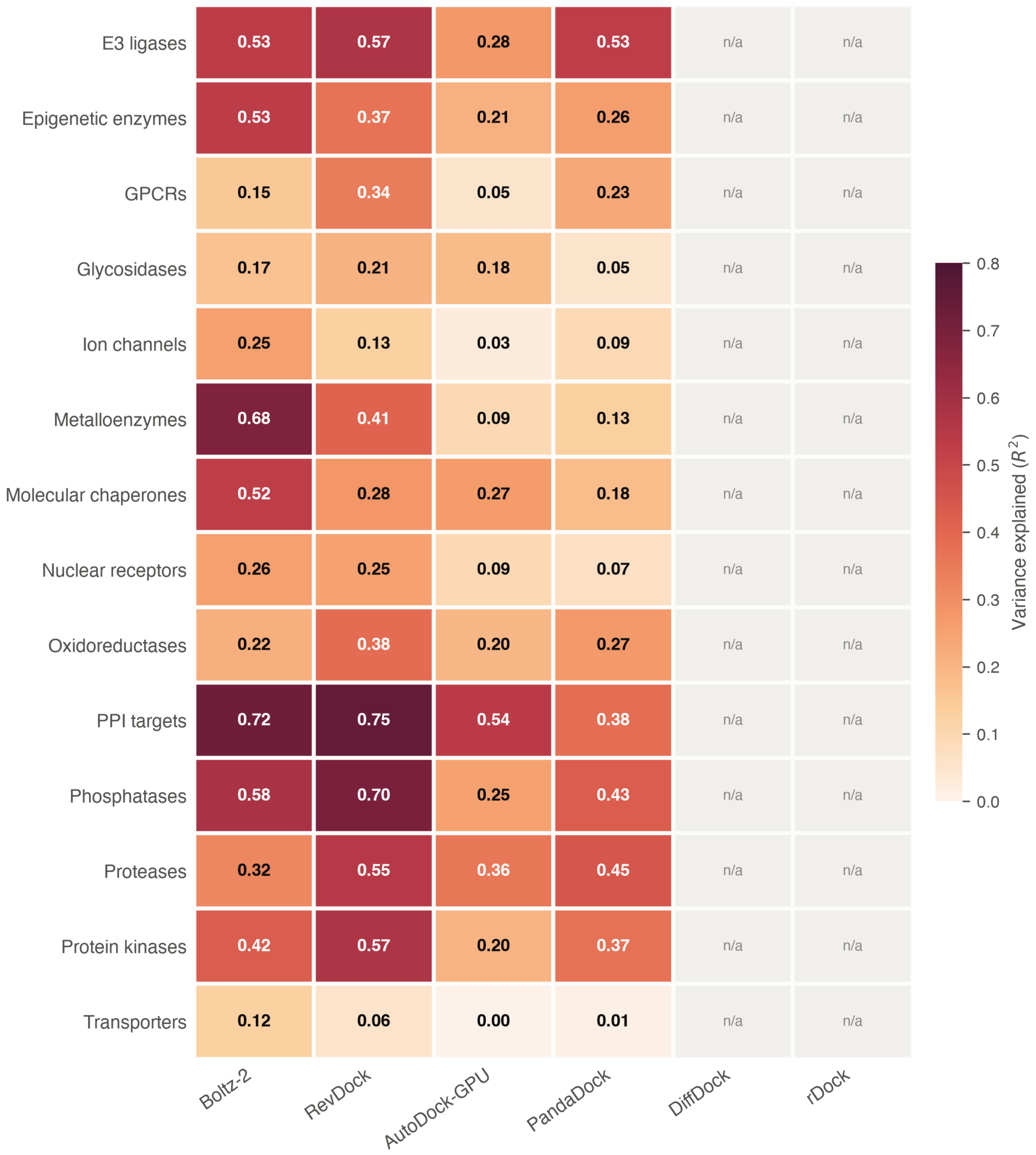


**Figure S14:** Scoring power by target family and engine ($R^2$), reproducing the main-text matrix with all six engines and with the two rank-only engines marked `n/a`.

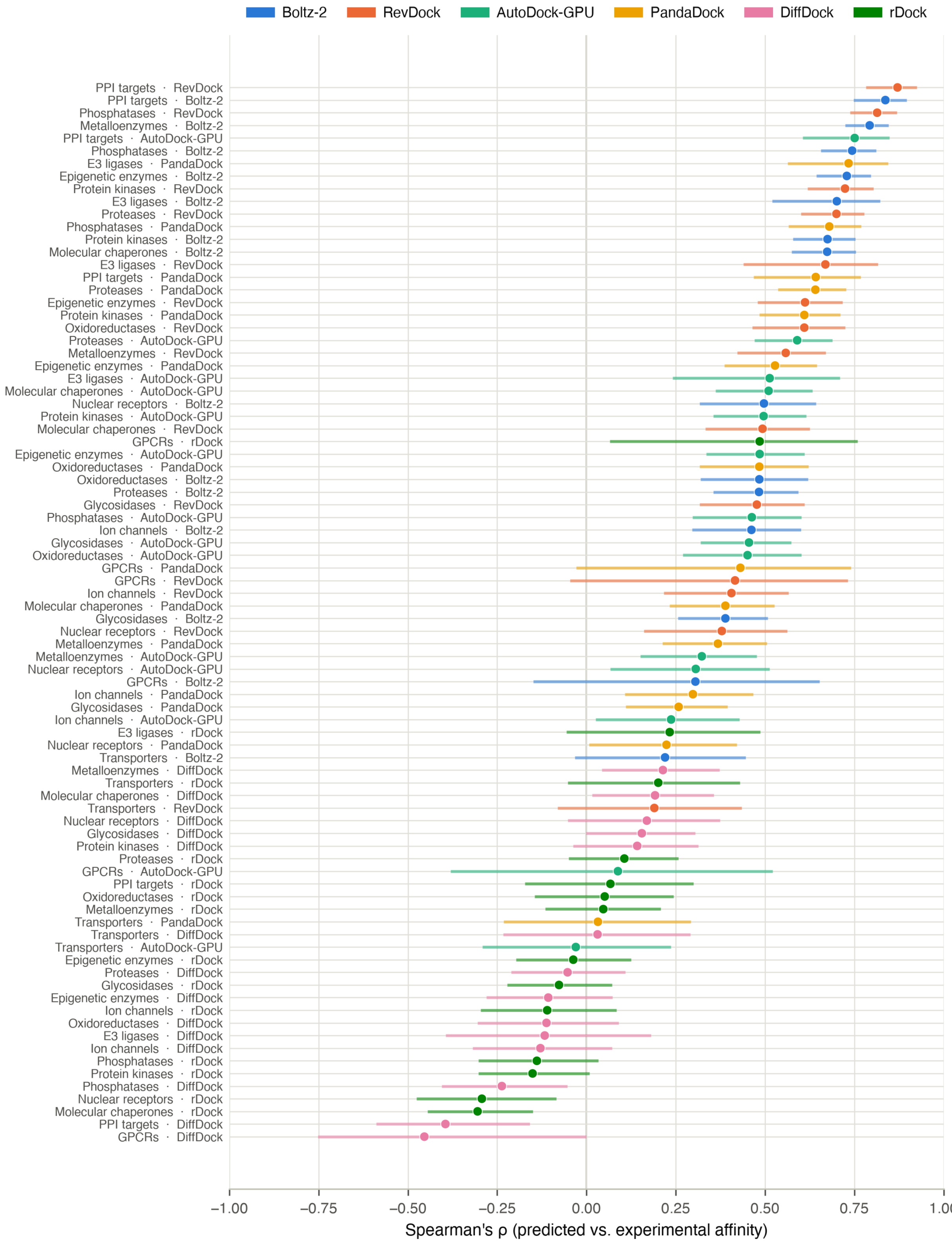


**Figure S15:** All engine × family pairs ranked by Spearman $\rho$ with 95% confidence intervals, sorted from weakest to strongest. An alternative presentation of Figure S7.

**Table S2:** Classification of the 14 protein families into six mechanistic groups based on which quantum mechanical simplification is most severely stressed.

| Group | Protein Families | Binding Site Character | Primary Limitations Stressed |
|---|---|---|---|
| **Membrane Embedded Proteins** | GPCRs, Ion Channels, Transporters | Proteins embedded within the lipid bilayer cycling between distinct functional states; GPCRs cycle between active and inactive states, ion channels open and close, and transporters adopt different conformations to transport molecules through the membrane | *Implicit solvent:* the dielectric constant within the membrane creates a large discrepancy relative to the aqueous assumption as a result of the hydrophobic nature of the lipid bilayer, rendering electrostatic scoring incorrect for membrane-embedded binding sites. *Rigid receptor:* conformational changes associated with biological function are not captured |
| **Metal Coordination Proteins** | Metalloenzymes, Epigenetic Enzymes (HDACs) | Metal ion embedded in the active site, typically zinc, coordinated by a tetrahedral shell of two Histidine residues, a Glutamate or Aspartate, and a displaced water molecule | *Pairwise additivity:* the interaction is not the sum of all interactions in the tetrahedral coordination shell; all interactions are dependent on one another. *Fixed point charges:* zinc is highly polarizable and the fixed charge cannot capture the electron density shift |
| **Deep Rigid Pocket Proteins** | Protein Kinases, Proteases, Nuclear Receptors, Phosphatases | Deep well-defined pockets in soluble proteins lacking metal ions; dominant interactions are hydrophobic contacts and discrete hydrogen bonds | All four limitations are least stressed; hydrophobic contacts and hydrogen bonds are approximately additive; the aqueous dielectric assumption is appropriate for soluble proteins |
| **Flexible and Shallow Pocket Proteins** | Molecular Chaperones, Oxidoreductases | Shallow pockets with dynamic boundaries; molecular chaperones have a flexible lid controlling access to the active site; oxidoreductases have a cofactor within or adjacent to the active site affecting pocket geometry | *Rigid receptor:* engines docking into a single fixed conformation will systematically fail for ligands that induce lid closure in molecular chaperones or require the cofactor-bound pocket geometry in oxidoreductases |
| **Large Flat Interface Proteins** | PPI Targets, E3 Ligases | Large flat protein-protein interfaces where binding energy is concentrated at a small number of hot spot residues; E3 ligases connect CRBN or VHL to a target protein via a PROTAC or Molecular Glue involving two binding events | *Pairwise additivity:* multiple simultaneous contacts are required between the inhibitor and residues; all interactions are dependent on one another. *Implicit solvent:* displacing ordered water molecules from a large flat interface is a cooperative thermodynamic process that pairwise implicit solvent terms cannot accurately model |
| **Polar Water-Dependent Proteins** | Glycosidases | Active site lined with polar and charged residues creating a highly ordered water network; unlike most hydrophobic binding sites, glycosidase active sites are entirely polar | *Implicit solvent:* water displacement thermodynamics cannot properly be modeled. *Fixed point charges:* polarization of charged residues in a purely polar environment is not captured. *Pairwise additivity:* each H-bond influences the geometry and strength of adjacent H-bonds through the water network |
